# Comparative chemical genomics in *Babesia* species identifies the alkaline phosphatase phoD as a novel determinant of resistance

**DOI:** 10.1101/2023.06.13.544849

**Authors:** Caroline D. Keroack, Brendan Elsworth, Jacob A. Tennessen, Aditya S. Paul, Renee Hua, Luz Ramirez-Ramirez, Sida Ye, Cristina M. Moreira, Marvin J. Meyers, Kourosh Zarringhalam, Manoj T. Duraisingh

**Affiliations:** Department of Immunology and Infectious Diseases, Harvard T.H. Chan School of Public Health, Boston, MA, USA; Department of Mathematics, University of Massachusetts Boston, Boston, MA, USA; Department of Chemistry, Saint Louis University, St. Louis, MO, USA; Center for Personalized Cancer Therapy, University of Massachusetts Boston, Boston, MA, USA

## Abstract

*Babesiosis* is an emerging zoonosis and widely distributed veterinary infection caused by 100+ species of *Babesia* parasites. The diversity of *Babesia* parasites, coupled with the lack of potent inhibitors necessitates the discovery of novel conserved druggable targets for the generation of broadly effective antibabesials. Here, we describe a comparative chemogenomics (CCG) pipeline for the identification of novel and conserved targets. CCG relies on parallel *in vitro* evolution of resistance in independent populations of evolutionarily-related *Babesia* spp. (*B. bovis* and *B. divergens*). We identified a potent antibabesial inhibitor from the Malaria Box, MMV019266. We were able to select for resistance to this compound in two species of *Babesia,* achieving 10-fold or greater resistance after ten weeks of intermittent selection. After sequencing of multiple independently derived lines in the two species, we identified mutations in a single conserved gene in both species: a membrane-bound metallodependent phosphatase (putatively named PhoD). In both species, the mutations were found in the phoD-like phosphatase domain, proximal to the predicted ligand binding site. Using reverse genetics, we validated that mutations in PhoD confer resistance to MMV019266. We have also demonstrated that PhoD localizes to the endomembrane system and partially with the apicoplast. Finally, conditional knockdown and constitutive overexpression of PhoD alter the sensitivity to MMV019266 in the parasite: overexpression of PhoD results in increased sensitivity to the compound, while knockdown increases resistance, suggesting PhoD is a resistance mechanism. Together, we have generated a robust pipeline for identification of resistance loci, and identified PhoD as a novel determinant of resistance in *Babesia* species.

**Highlights:**

- Use of two species for *in vitro* evolution identifies a high confidence locus associated with resistance
- Resistance mutation in phoD was validated using reverse genetics in *B. divergens*
- Perturbation of phoD using function genetics results in changes in the level of resistance to MMV019266
- Epitope tagging reveals localization to the ER/apicoplast, a conserved localization with a similar protein in diatoms
- Together, phoD is a novel resistance determinant in multiple *Babesia spp*.

## Introduction

Babesiosis is a febrile illness caused by the tickborne apicomplexan parasite genus *Babesia. Babesia* has long been recognized as a disease of tremendous veterinary and agriculture importance, causing hundreds of millions of dollars of economic losses every year [1,2]. Babesiosis has steadily been gaining recognition as an important human zoonotic disease [3–6]. While usually vector borne, human infections can also be acquired through blood transfusion [7–14]. Symptoms are generally most severe in splenectomized or immunocompromised patients and the elderly [5,6,15]. Immunocompromised patients often experience treatment failure, occasionally leading to death [4,16]. Further, the true burden of disease remains unknown-several recent serological studies suggest far more widespread infection than reported [7,17–19].

There is a vast diversity of parasites, capable of infecting nearly any vertebrate host, with at least 100 described species [6,20,21]. This poses threats to wild life, companion animals, livestock, and provides a seemingly endless source of zoonotic infection. Bovine babesiosis caused mainly by *B. bovis* and *B. bigemina* accounts for significant agricultural losses globally [2,22–25]. While *B. microti* is the most common human pathogen in the USA, human babesiosis is caused by a range of parasites world-wide including *B. duncani*, *B. venatorum, B. crassa, B. divergens* and related parasites [5,6,26–29]. Of these, *B. divergens,* while rare, is often fatal [27,30]. The vast diversity of *Babesia* parasites, in addition to host plasticity, demands the development of a species-transcendent antibabesial drug which targets core, conserved biology.

The first line treatment for babesiosis in humans consists of atovaquone and azithromycin, while second line treatment for severe, refractory, or relapsing *Babesia* is a combination of quinine and clindamycin [15,31]. *De novo* resistance has been reported in patients, resulting in life threatening illness and even death [4,15]. The major treatments for veterinary *Babesiosis* are imidocarb dipropionate, diminazene aceturate, as well as an array of antibiotics such as doxycycline and clindamycin. While effective, both drugs suffer from major toxic side effects [24]. Highly variable efficacy of treatments between species is another major pitfall of current therapeutic strategies. For example, there is a 50-100 fold range in efficacy of atovaquone between the three major species known to infect humans-*B, divergens, B. microti,* and *B. duncani-*highlighting the need to identify a novel therapeutic strategy with efficacy across species [32–34]. Furhter, the reported efficacy of imidocarb ranges from an effective dose of ∼0.0001 μg/mL in *B. bovis* to ∼0.01 μg/mL in *B. divergens* to completely ineffective in *B. gibsoni* [35–37]. Conversely, the efficacy of diminazene aceturate is fairly consistent across species, however drug toxicity remains a major issue for therapeutic use [36,38,39]. Taken together, these pitfalls of current antibabesial therapy strongly warrant investigations into novel treatments.

Leveraging antimalarials for antibabesial development is strategic and cost effective. The Malaria Box compound library has been made publicly available by the Medicines for Malaria Ventures (MMV) [40], and studies that many of these compounds are effective against *Babesia* [34,40–42]. The structural diversity of the library furthermore allows for deep exploration of parasite targets and cellular phenotypes, and metabolomics suggests these compounds target many distinct pathways [40,43]. With the highly shared chemosensitivity, the Malaria Box provided an ideal resource for identifying novel, species transcendent targets in *Babesia*.

The majority of previous efforts to identify novel target-inhibitor pairs in apicomplexans using *in vitro* evolution and chemical genomics have focused on single species [44,45]. Many target-inhibitor pairs have been identified using *in vitro* resistance evolution and chemical genomics in *Plasmodium falciparum*, including those in the malaria box, suggesting the utility of this method in rapidly identifying target-inhibitor pairs [44–51]. However, performing experiments in parallel species allows the opportunity to rapidly identify conserved resistance mechanisms and druggable targets (**Fig 1A**). Indeed, a comparative study of *in vitro* evolution between *P. falciparum* and *Saccharomyces cerevisea* identified a novel target-inhibitor pair with high confidence [52]. This underscores the utility of taking a comparative approach to target discovery. While the major etiological agent of human babesiosis, *B. microti*, cannot be cultured *in vitro,* dozens of other species are readily cultured *in vitro*, including *B. bovis* and *B. divergens* [20,53,54]. Previous studies of drug resistance in *Babesia* have relied on *in vitro* evolution[37,39,55,56], however to date, none have paired this technique with full genome sequencing to identify genetic loci associated with resistance. Further, both *B. bovis* and *B. divergens* are amenable to genetic manipulation [57–59]. Here, we use these two species to develop the CCG pipeline, and identify a novel, conserved resistance mechanism.

**Figure 1.**
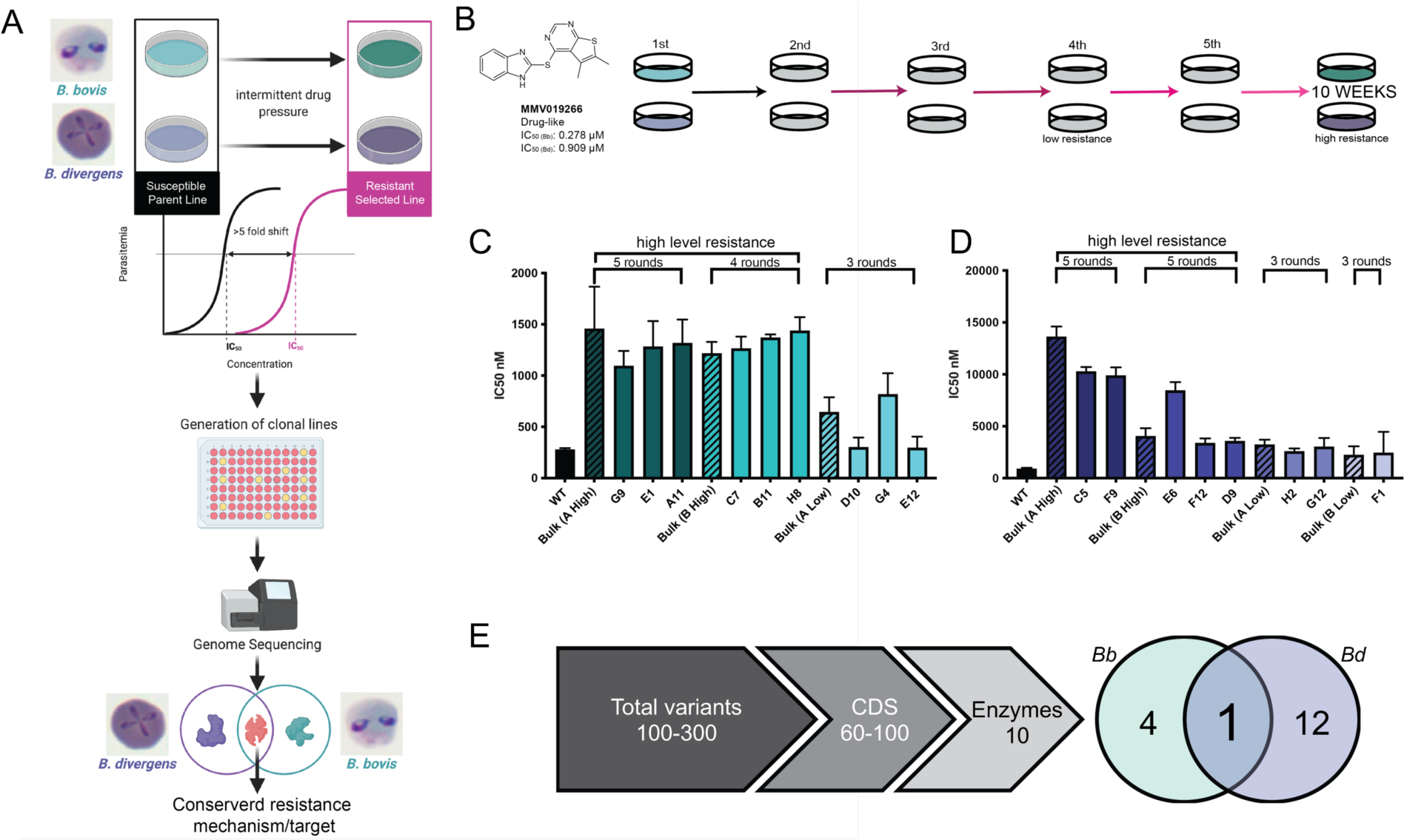
Comparative chemical identifies a single locus associated with high level resistance to MMV019266. (A) Schematic representation of the comparative chemical genomics pipeline (CCG). Representative light microscopy images *B. bovis* is outlined in teal, *B. divergens* is outlined in purple. The pipeline begins with in vitro selection for resistance in two species, followed by whole genome sequencing, and validation of potential gene candidates. (B) The structure of MMV109266 and IC_50_ values is next to a schematic of intermittent drug selection. Low level resistance was achieved by three rounds of selection, and high by five rounds. Selections were given a maximum of 10 weeks to yield resistant parasite populations of both species for this study. For each species, selections were run in duplicate. (C) *B. bovis*: Successful resistance selection in both species to MMV019266. High level resistance was achieved after 5 rounds of selection. Hashed bars represent the IC_50_ of the bulk population, solid bars represent the IC_50_ of clonal lines derived from the bulk population. Error bars represent standard deviation between 3 independent dose-response curves. (D) *B. divergens* resistant clones. High level resistance was achieved after 5 rounds of selection. Hashed bars represent the IC_50_ of the bulk population, solid bars represent the IC_50_ of clonal lines derived from the bulk population. Error bars represent standard deviation between 3 independent dose-response curves. (E) Whole genome sequencing identified a single conserved gene associated with resistance to MMV019266 after filtering steps to remove non CDS mutation, mutations in variable surface antigens, and select for mutations in genes with predicted enzymatic domains.

Using the previous screening of the Malaria Box in *B. divergens* and *B. bovis* as a resource [34], we selected a top candidate compound for target identification (MMV019266) from this screen based on its similar potency in both *B. bovis* and *B. divergens* and the compound having a drug like structure. We used *in vitro* evolution select for resistance, followed by whole genome sequencing of resistant clones to identify candidates associated with compound resistance. Though this, we identified a single, species transcendent locus associated with MMV019266 resistance. We subsequently used recently developed genetic methods to validate and investigate the function of this previously uncharacterized membrane-bound metallodependent phosphatase, PhoD [57]. Such phosphatases have been studied deeply in prokaryotes [60–64]. However, in eukaryotes they have been only investigated and localized in marine algae [65–67], which are distant relatives of apicomplexans (chromalveolates). In all systems studied these PhoD alkaline phosphatases play important roles in phosphate starvation through salvaging of inorganic phosphate either from the environment or from internal stores [64,68]. Using functional genetics, we investigated the relationship between MMV019266 and PhoD. We demonstrate that this protein localizes to the apicoplast and endoplasmic reticulum (ER), which are observed locations in the marine algae [65,66].

## Methods

### Parasite culture

The *Babesia bovis* strain MO7 provided by David Allred of the University of Florida was maintained in purified bovine RBCs (hemostat) at 4% hematocrit in RPMI-1640 media supplemented with 25 mM HEPES, 11.50 mg/l hypoxanthine, 2.42 mM sodium bicarbonate, and 4.31 mg/ml AlbuMAX II (Invitrogen). Before addition of AlbuMAX-II and sodium bicarbonate, we adjusted the pH of the media to 6.75. *Babesia divergens* strain Rouen 1987, kindly provided by Kirk Deitsch and Laura Kirkman (Weill Cornell Medical College), was maintained under the same conditions in purified Caucasian male O+ human RBCs (Research Blood components). All cultures were maintained at 37 °C in a hypoxic environment (1% O_2_, 5% CO_2_). Clonal lines of parasites were used for all selections and were derived from the provided strains via limiting dilution-these will be referenced as BdC9 (*B. divergens*) and BOV2C (*B. bovis*).

### Compounds and reagents

MMV0190266 (Vitascreen, LLC), Atovaquone (Sigma Aldrich Cat. No. PHR1591), Imidocarb dipropionate (Sigma Aldrich Cat. No. 33441), WR99210 (Jacobus Pharmaceuticals) were prepared in DMSO. Clindamycin (Sigma Aldrich Cat. No. 1136002), azithromycin (Sigma aldrich Cat. No. 1046056), diminazene aceturate (Sigma Aldrich Cat. No. D7770), puromycin (Sigma Aldrich Cat. No. P8833), and blasticidin S (Invivogen Cat. No. ant-bl-10p) were prepared in water. Isopentanyl pyrophosphate (IPP, Isoprenoids LLC), and geranylgeraniol (Sigma Aldrich, Cat. No. G3278) were prepared in ethanol. Compound-1 (DMSO-based) was a gift from Dr. Jeffrey Dvorin (Boston Children’s Hospital). Shld1 was synthesized and prepared as previously described [69,70]. Sources and dilutions for antibodies used in immunoblotting and immunofluorescence analysis are as follows: chicken α-GFP (IFA-1:500, abcam ab13970), rabbit α-BIP was a gift from Dr. Jeffrey Dvorin (IFA-1:500, Boston Children’s Hospital), rat α-HA 3F10 (WB 1:1000, Roche Cat. No. 11867423001), rabbit α-H3 (WB 1:8000, abcam Cat No. ab1791), rabbit α-mCherry (IFA 1:1000, abcam Cat. No. ab183628). The secondary antibodies used for IFA were Alexa-Fluor 488 and 594-conjugated antibodies against chicken or rabbit IgG diluted as recommended by the manufacturer (Invitrogen Cat. Nos. chicken 488 A21207, rabbit 594 A11039). Secondary antibodies for immunoblotting were IRDye® 680 LT against rat, or rabbit IgG diluted per manufacturer’s instructions (LICOR/Odyssey Cat. Nos. rat 926-68029, rabbit 925-68023).

### Plasmids

Primers for PCR amplification and verification genetic manipulation in *B. divergens* are shown in the **SI table 1**. The plasmids for generation of transgenic parasites containing mutations in *phoD* were built using pBdEF-Cas9-BSD as a base, and will be known subsequently as pBdEF-Cas9-BSD-phodR [57]. To generate pBdEF-Cas9-BSD-phodR a 500 base pair homology repair template, designed to the 3’ end of the gene, with the desired resistance mutation (444G>C) and a silent Cas9 shield mutation (424G>A) was constructed by PCR sewing and inserted between sites AvrII and PacI. The resulting plasmid was further modified with the guide RNA via insertion at the BbsI restriction site, resulting in pBdEF-Cas9-BSD-phodR-g1. This results in a single plasmid strategy for gene editing. Homology fragments were assembled using the NEBuilder® 2 X HiFi DNA assembly master mix (NEB, Cat. No. #E2621), while guide RNAs were inserted by ligation using T4 ligase (NEB, Cat. No. M0202). Plasmids generated for end-tagging of the endogenous locus were also based on pBdEF-Cas9-BSD. For generation of the homology repair, the 3’ end of *phoD* (250 bp) was amplified without the stop codon intact followed by the tag, either HA-DD-glmS (inducible knockdown), GFP with a 3’ stop codon, and the 3’ UTR (250 bp) of *phoD* were all PCR amplified, primers can be found in **SI table 1**. The GFP tag was preceded by a flexible linker sequence GGGGS. Upon generation, fragments were sewn into a signal fragment by PCR. Fragments were assembled using the NEBuilder® 2 X HiFi DNA assembly master mix (NEB, Cat. No. #E2621). The homology and guide RNA were inserted as described for pBdEF-Cas9-BSD-phodR, and resulted in plasmids pBdEF-Cas9-BSD-phod-DD and pBdEF-Cas9-BSD-phodGFP. All guide sequences are available in **SI table 1**. Plasmids were validated by sanger sequencing.

Plasmids generated for episomal expression (including localization and over expression) were generated via insertion of various synthesized fragments into pBdEF-GFP-BSD using restriction sites XhoI and SpeI. All synthesis products were made by Twist biosciences. The synthesis products SP+TP_lytB_-(mCherry or GFP) were generated by fusing the first 213 nucleotides of lytB mRNA (predicted signal and transit peptide) to the fluorescence protein sequence (mCherry or GFP) [71]. Similarly, SP+TP_phoD_-GFP was produced by adding the first 210 nucleotides of the *phoD* mRNA. The resulting plasmids will be referred to as pBdEF-SP+TP_phoD_-GFP-BSD, pBdEF-SP+TP_lytB_-GFP-BSD, pBdEF-SP+TP_lytB_-mCherry-BSD, pBdEF-SP_lytB_-mCherry-BSD. Cytoplasmic mCherry expression was done by swapping the fluorescent marker in the base plasmid to form pBdEF-mCherry-BSD. The overexpression plasmid was prepared by inserting the gDNA sequence of *phoD* generated by PCR on genomic DNA, as well as swapping the Bdiv_030590/Bdiv_030580c bidirectional promoter to the endogenous phoD promoter to drive phoD expression, and the Bdiv_021550 promoter driving BSD. The reverse primer excluded the stop codon and contained a 3XHA tag (followed by a stop), resulting in HA tagged fragments. Fragments were assembled using the NEBuilder® 2 X HiFi DNA assembly master mix (NEB, Cat. No. #E2621). The resulting plasmid will be referred to as pBd-phoD-HA-BSD.

### Synchronous phenotyping assays

Parasites were synchronized by mechanical release from the red blood cell as previously described [30,72]. In brief: 30 mL of culture (4% HCT, 1.2 mL packed RBCs) was grown to high parasitemia (>20%), centrifuged and resuspended in 10 mL of complete media. For *B. divergens,* the suspension was passed through a 1.2 μm syringe filter; *B. bovis* was passed first through a 5 μm syringe filter followed by a 2 μm syringe filter. After mechanical release, the parasite suspension was centrifuged at 3000 × G for 5 min to pellet the merozoites, then added to 1 mL of 20% hematocrit blood in complete media and allowed to reinvade for 30 m at 37 °C, shaking at 600 rpm. Reinvaded RBCs were then washed, and finally put in culture supplemented with 50 μg/mL of heparin sulfate to prevent reinvasion. Immediately after reinvasion, parasites were put on MMV019266 continuously at 10 X IC_50_. Samples were taken every 4 hours for 12 hours and fixed in 4% PFA + 0.025% glutaraldehyde in 1 X PBS overnight at 4 C. Samples were then prepared for flow cytometry to access nuclear content. To assess nuclear content, iRBCs were prepared for flow cytometry via fixation in a solution of paraformaldehyde (%) and glutaraldehyde (%) in a 1 X PBS. Fixed cells were maintained at 4C until the end of the time course experiment, then washed with 1 X PBS, permeabilized with triton 100 (%), treated with RNase A (amount), and finally stained with SYBR green (1: 10,000). Data were processed using FloJo, and downstream analysis was performed using GraphPad Prism v9 (GraphPad Software, Inc).

### Dose-response assays in *B. bovis* and *B. divergens*

The assays to determine the IC_50_ of all compounds were done in 96 well plates as previously described in *B. divergens* [73,74], with several modifications. Asynchronous cultures were prepared at 0.5% parasitemia in 3% hematocrit, 50 μL were added to wells of a black, clear bottom (costar) 96 well plate. For apicoplast rescue experiments, the media was supplemented with either 300 μM IPP or 5 μM geranylgeraniol. Compounds were prepared by serial dilution. Negative controls included no drug and 1 % DMSO. Puromycin was included as killing (positive) control. Parasites were then incubated with compounds for 72 hours in standard culture conditions. Subsequently, cultures were lysed with a solution containing SYBR green (1:5000, Invitrogen Cat. No. S7567) (Lysis buffer: 0.16% saponin (Calbiochem/EMD #558255), 0.05 M TRIS-HCl (Roche Cat. No. 10812846001), 0.8 mM EDTA (Sigma Aldrich Cat No. E7889), 1.6% Triton X-100 (Sigma Aldrich Cat. No. T8787). Sybr green fluorescence was measured on a SpectraMax® iD5 (Molecular Devices) and DNA content is used as a proxy for parasite growth.

### *In vitro* resistance generation

Rounds of intermittent drug selection were performed to attain resistance to MMV019266 in both *B. divergens* and *B. bovis.* Parasites were exposed to 5 × IC_50_ for 3-5 days until growth was halted and parasites appeared punctate. Parasites were allowed to recover until normal growth resumed (between 2-3 weeks for each round of selection). Parasites were subjected to rounds of selection until a greater than or equal to 5 fold shift in IC_50_ was achieved. Clonal lines of resistant parasites were then generated by limiting dilution.

### Whole genome sequencing

Genomic DNA of parasites was isolated using the Qiagen DNeasy blood and tissue kit, following manufacturer’s guidelines (Qiagen Cat. No. 69504). Clonal lines of *B. bovis* and *B. divergens* resistant to MMV019266, as well as the starting parental clonal lines, were prepared for sequencing using the Illumina Nextera XT workflow. Libraries were pooled (8pM) and sequenced on the Illumina MiSeq using v2 reagents (500 cycle, paired end). FastQ files were assessed for quality using FastQC and low quality bases were trimmed used trimmomatic [75]. All data are publicly available and have been deposited in the SRA under the accession number: PRJNA924801.

### Identification and confirmation of single nucleotide polymorphisms and copy number variation (CNV) analysis

Cleaned, paired fastq files were aligned to the respective reference genome (*B. divergens* strain 1802A [76], *B. bovis* T2Bo [77,78] both accessed via piroplasmaDB) using BWA [79]. Alignments were sorted and subsequently merged into VCF files using SamTools [80], BedTools [81], and VCFtools [82]. SNPs were filtered first by comparison to the sequenced parental clone (divergens-BdC9, bovis-BOV2C) – rows were retained if any clone contained a variant in comparison to the parental strain. Variants were then filtered by quality score (>30), and heterozygous calls were removed (*Babesia* is a haploid organism). Next, variants were filtered based on location in coding sequences, and further refined based on functional annotation-all variants in antigen superfamilies were removed, as well as most hypothetical or unannotated genes, retaining variants in only genes with predicted enzymatic function. SNPs were then mapped to their respective sequences and assessed for functional significance (missense, nonsense, etc). Finally, the candidate gene sets were overlapped in *B. bovis* and *B. divergens* to identify possible conserved resistance determinants. To evaluate the role of copy number variants in MMV019266 resistance, read counts were evaluated from the alignment (BAM) files to evaluate sequence depth across the genome.

### N-ethyl-N-nitrosourea mutagenesis

Parasites were grown to 5% parasitemia in 10 mL of culture at 4% hematocrit. Parasites were then exposed to either 1 mM or 1.5 mM N-ethyl-N-nitrosourea (ENU) for 4 hours. ENU was then washed off and parasites were allowed to recover, until growth returned to 5% parasitmia after death induced by ENU. Upon recovery, parasites were placed on 10 X IC_50_ of MMV019266 or atovaquone (control). Parasitemia was monitored daily until robust growth was observed. Parasites were assessed for mutations in the gene of interest by PCR.

### Transfection of parasites

Free merozoites, isolated as described, were used for amaxa transfection as previously described [57]. Briefly, the pellet containing free merozoites and cell debris, was resuspended in 110 µl P3+DNA solution (Lonza Cat. No. V4Xp-3024). Transfection of either free merozoites carried out in a 4D-Nucleofector System (Lonza), using the FP158 electroporation settings. After electroporation of free merozoites, the parasite and buffer mixture was immediately transferred to 1 ml of RPMI containing 200 µl packed RBCs and pre-heated to 37 °C. Parasites were allowed to invade at 37 °C shaking at 600 rpm for 30 mins before being washed with 10 ml RPMI to remove the P3 solution and returned to culture. Parasites were returned to culture at a final volume of 10 mL RPMI at 2% HCT. Parasites were allowed to recover for 24 hours prior to addition of blasticidin.

### Protein sequence alignments

All apicomplexan sequences were downloaded from piroplasmaDB, toxoDB, and plasmoDB (**SI Table 2**). To identify genes containing phoD domains in apicomplexa, we performed a search of these databases for the InterPro domain IPR018946 (“Alk_phosphatase_PhoD-like Alkaline phosphatase D-related” and “PhoD-like_MPP PhoD-like phosphatase, metallophosphatase domain”) as well as BLAST of the phoD domain of BdPhoD. *Bacillus subtilis* sequence was accessed from NCBI genbank (Bsubtilis_AAB47803). *Phaeodactylum tricornutum* phoD sequences were acquired from the Phatr2 database (PtPhos3_45959, PtPhos4_39432, PtPhos5_48970, PtPhos6_45757, PhPhos7_45174). PhoD domains were extracted from full sequences based on the PFAM predictions for each protein. Alignments of whole protein sequences and domain sequences were performed using Geneious v9.1.5, with standard alignment parameters (Neighbor joining clustering method, ClustalW alignment). Phylogenetic trees were constructed using Geneious v9.1.5, using Jukes-Cantor genetic distance model, the UPGMA tree build model, based on a global alignment with a Blosum62 cost matrix.

### Protein structure modeling and molecular docking

The structure of PhoD was predicted using Alphafold [83]. Protein structure alignments were performed in PyMol v2.3.2. Docking studies were conducted with Schrodinger Maestro Release 2021-1 and 2022-3. The crystal structure for *Bacillus subtilis* PhoD (2YEQ) was imported from the Protein Data Bank (PDB). Both the model *B. divergens* PhoD and *B. subtilis* PhoD structures were prepared using Protein Preparation Wizard and aligned. Glide docking grids were generated for the active site in these structures, placing no restrictions or constraints on the ligand binding protocol. The ligand MMV019266 was prepared using LigPrep. Glide docking was performed using these Glide docking grids and MMV019266 under the XP precision mode and all other defaults with the exception of increasing the number of output structures to three per docked compound. SiteMap was utilized to identify five potential binding sites in the *B. divergens* PhoD models.

### Immunoblotting of parasite lysates

Samples were taken for comparison at the same parasitemia. Infected red blood cells were pelleted, then lysed with a solution of 0.015 % saponin in 1 X PBS with protease inhibitors (Millipore Sigma Cat. No. 11873580001) Parasites were pelleted and dissolved in sample buffer (Cell Signaling Technologies Cat. No. 7722) prepared via manufacturer’s instructions. Samples were denatured at 95 °C for 3 minutes prior to loading onto a 16.5% Mini-PROTEAN® Tris-Tricine Gel (Biorad Cat. No. 4563066). Gels were transferred to a nitrocellulose membrane (Millipore Sigma Cat. No. GE10600002) in cold wet transfer buffer overnight at 15 V, followed by 1 hour at 40 V. Membranes were blocked in Intercept (PBS) Blocking Buffer (LI-COR Cat. No. 927-70001) then treated with primary followed by secondary antibodies. Bound antibodies were detected using IR Dyes on the LiCor Odyssey Clx imager. Band intensity was quantified using Fiji version 2.3.0.

### Validation of causal mutations by reverse genetics

To introduce the single nucleotide polymorphism to generate a C196W mutation in PhoD in *B. divergens,* parasites were transfected with pBdEF-Cas9-BSD-phodR. Parasites were exposed to blasticidin 24 hours post transfection, and were kept on blasticidin for 3 days. Parasites were allowed to recover until recrudescence. Gene editing was validated by PCR of PhoD with primers that were designed outside of the homology region in the vector (**SI table 1**). Parasites were cloned by limiting dilution, and assessed for gene editing. Edited clones were then tested for resistance to MMV019266 as described.

### Overexpression of BdPhoD

Overexpression of *phoD* was achieved via transfection of pBd-phoD_gDNA_HA-BSD. Parasites were maintained on blasticidin 24 hours after transfection. Expression of phoD-HA was assessed by immunoblotting. Parasite proliferation was monitored by thin blood smear and flow cytometry (sybr green staining) over 72 hours starting from 0.1 % parasitemia, in comparison to non-transfected wild-type parasites.

### Conditional knockdown of BdPhoD

Conditional knockdown of PhoD was achieved by transfecting parasites with pBdEF-Cas9-BSD-phod-DD. Parasites were maintained on blasticidin 24 hours after transfection until no parasites were observed-at which time blasticidin was removed. Parasites were then maintained on 500 nM Shld1. Tagging of *phoD* was confirmed by PCR (**SI Fig 5, SI table 1**). Parasite proliferation +/- Shld1 was monitored by thin blood smear and flow cytometry (sybr green staining) over 72 hours starting at 0.1% parasitemia. This was also compared to wild type growth rate. Effect of Shld1 titration on tagged parasite lines was assessed using the fluorescence based IC_50_ assay previously described. Degree of knockdown was assessed by immunoblotting on a nitrocellulose membrane using α-HA, normalized to α-H3. To determine IC_50_ with MMV019266, Shld1 was washed (4X) off the parasites immediately prior to set up of the assay. Parasites were exposed to compound for 72 hours prior to lysis as previously described.

### Knock out strategy for phoD

To attempt to knock out phoD we used a CRISPR/Cas9 approach. Parasites were co-transfected with linear PCR product (repair template synthesized by Twist Bioscience, San Fransisco, CA) containing the BSD resistance cassette flanked by 500 bases of homology in the 5’ and 3’ end of phoD, and two plasmids containing gRNA targeting the 5’ and 3’ ends of phoD. The gRNA plasmids were generated from the base plasmid pBdEF-Cas9-BSD. The resulting plasmid was further modified with the guide RNA via insertion at the BbsI restriction site, resulting in pBdEF-Cas9-BSD-phodKO-g1 and pBdEF-Cas9-BSD-phodKO-g2. Parasites were continuously maintained on blasticidin 24 hours after transfection. Clonal lines were generated by limiting dilution, and assessed for integration of the knock-out cassette and loss of phoD by PCR. All primers can be found in **SI table 1**.

### Localization of BdPhoD

Immunofluoresence assays (IFA) were performed as previously described [57,84– 86]. Briefly, thin smears of parasitized erythrocytes were air dried and fixed in ice-cold methanol, then permeabilized. For endoplasmic reticulum colocalization: chicken α-GFP was used to detect PhoD-GFP and was co-stained with rabbit α-BIP (ER). After antibody treatment, slides were stained with Hoechst 33342 (1:10000 Thermo Fisher Scientific Cat. No. H3570) for 2 minutes. For localization of the mitochondrion, live parasites (WT or GFP) were incubated with 400 nM MitoTracker**^TM^** Red CM-H2XRos (Invitrogen, #M7513) for 20 minutes at 37 °C then immobilized on slides coated with poly-L lysine. Parasites were pelleted and washed once with 1 X PBS. Parasites were then incubated with Hoechst 33342 (Thermo Fisher Scientific) for 5 minutes at RT. Parasites were dispensed onto poly-L-lysine coated slides to immobilize them for live cell imaging. Localization of the apicoplast was done by transfecting endogenously tagged parasites, phoD-GFP with SP+TP_lytB_-mCherry-BSD. All images were acquired on Zeiss Axio Observer using a 100x oil immersion lens. Analysis and quantification was all performed in FIJI version 2.3.0.

## Results

### Selection for resistance against MMV019266

We selected MMV019266 from a prioritized list of 10 most potent compounds from our previous screening of the malaria box to initiate the comparative chemical genomics pipeline [40,44]. This compound was selected of these 10 due to the similar potency in both *Babesia* species, the lack of a known target, and for being the only of the compounds with drug-like properties [34]. In both species, the IC_50_ was in the nanomolar range: 278.9 ± 13 nM in *B.bovis* and 909.1 ± 87 nM in *B. divergens. In vitro* evolution experiments were performed in parallel in two independent cultures of *B. divergens* and *B. bovis* for 10 weeks, with selections consisting of exposure to compound (5 × IC_50_) until no parasites were detectable (2-5 days) followed by a recovery period until recrudescence (1-2 weeks) (**Fig 1B**). In both *B. bovis* and *B. divergens* we were able to achieve at least a five-fold shift in IC_50_ to MMV019266 (**Fig 1 C,D**). Resistance was generated in a stepwise fashion, resulting in low and high resistance. Three rounds of intermittent drug selection in *B. bovis* resulted in a three-fold shift; after five rounds of selection, a seven-fold shift in IC_50_ (**Fig 1 C**). In *B. divergens* we observed a 2-3 fold shift in IC_50_ after 3 rounds of selection, and after five rounds of selection observed >10-fold shifts in IC_50_ in both independent selections (**Fig 1 D**). The IC_50_ values for all selections can be found in **SI Table 3**.

### Whole genome sequencing of resistant clonal parasites reveals a single conserved locus

For *B. bovis* we performed whole genome sequencing (WGS) of nine resistant clones from the two independent selections, and seven clones from the *B. divergens* selections (information about sequencing can be found in the (**SI Table 4**)). We sequenced clones with both low level and high level of resistance. We also sequenced the parental susceptible starting clone for comparison. Sequencing of *B. bovis* resulted in an estimated average coverage of 129-fold across the 9.4 Mb genome. The average estimated coverage of sequenced clones in *B. divergens* was 152.4-fold across the 10 Mb genome (**SI Table 4**). After alignment and filtering the mean depth per site in the genome ranged from 1.9-30.3 for *B. divergens*, and 5 – 21.8 for *B. bovis* (**SI Table 4**). To identify potential genetic changes in resistant clones, we generated VCF files and filtered variants in which at least one clone was different from the parental and reference strain. Based on studies in *Plasmodium falciparum* we reasoned that mutations of interest would be in the coding genome, and would most likely be in predicted enzymes [44]. Subsequently, we filtered the resulting SNP mutations identified across clones and species to include only those within CDS. We also reasoned that antigenic surface proteins and antigens would have inherently high variability [76], and removed these genes from the analysis. Finally, we excluded hypothetical proteins with no known annotations or homologous domains to related parasites (i.e. *Plasmodium falciparum*) (**Fig 1E**). In *B. bovis* and *B. divergens* resistant to MMV019266 across the clones we identified 5 and 13 possible gene candidates, respectively (**Fig 1E**). Of these, only one gene candidate was identified as mutated in both species: Bdiv_001570c and the orthologous BBOV_I003300, here after referred to as BdPhoD and BbPhoD respectively (**Fig 1E**). While Bdiv_0015750c is the only predicted phoD-like phosphatase in *B.* divergens, there appears to be a duplication of phoD in *B. bovis* (BBOV_I003305), however this gene is not mutated in the MMV019266 resistant parasite, and is less similar to BdPhoD than BBOV_I003300 (BbPhoD). In the WGS *B. divergens* samples, we identified a single amino acid that was mutated: C196W or C196F (**Fig 2A**). These mutations are located in the predicted phoD-like phosphatase domain of the gene. In *B. bovis* we identified three different mutations: G229D in the phoD-like phosphatase domain, and R418L/R418P in the C-terminal helix (**Fig 2A**). After genome sequencing identified phoD as a potential resistance candidate, we sequenced BbPhoD in two additional *B. bovis* resistant clones not used for WGS, and identified an additional mutation in clone B6 (G222W) (**Fig 2 A**). Mutations in phoD were only identified in the more highly resistant parasite clones, suggesting low level resistance occurs through an alternative mechanism that could not be identified through sequencing. We found no compelling evidence that copy number variation contributes to resistance to MMV019266 in either lowly or highly resistant parasites (information about CNV analysis can be found in **SI Information**).

**Figure 2.**
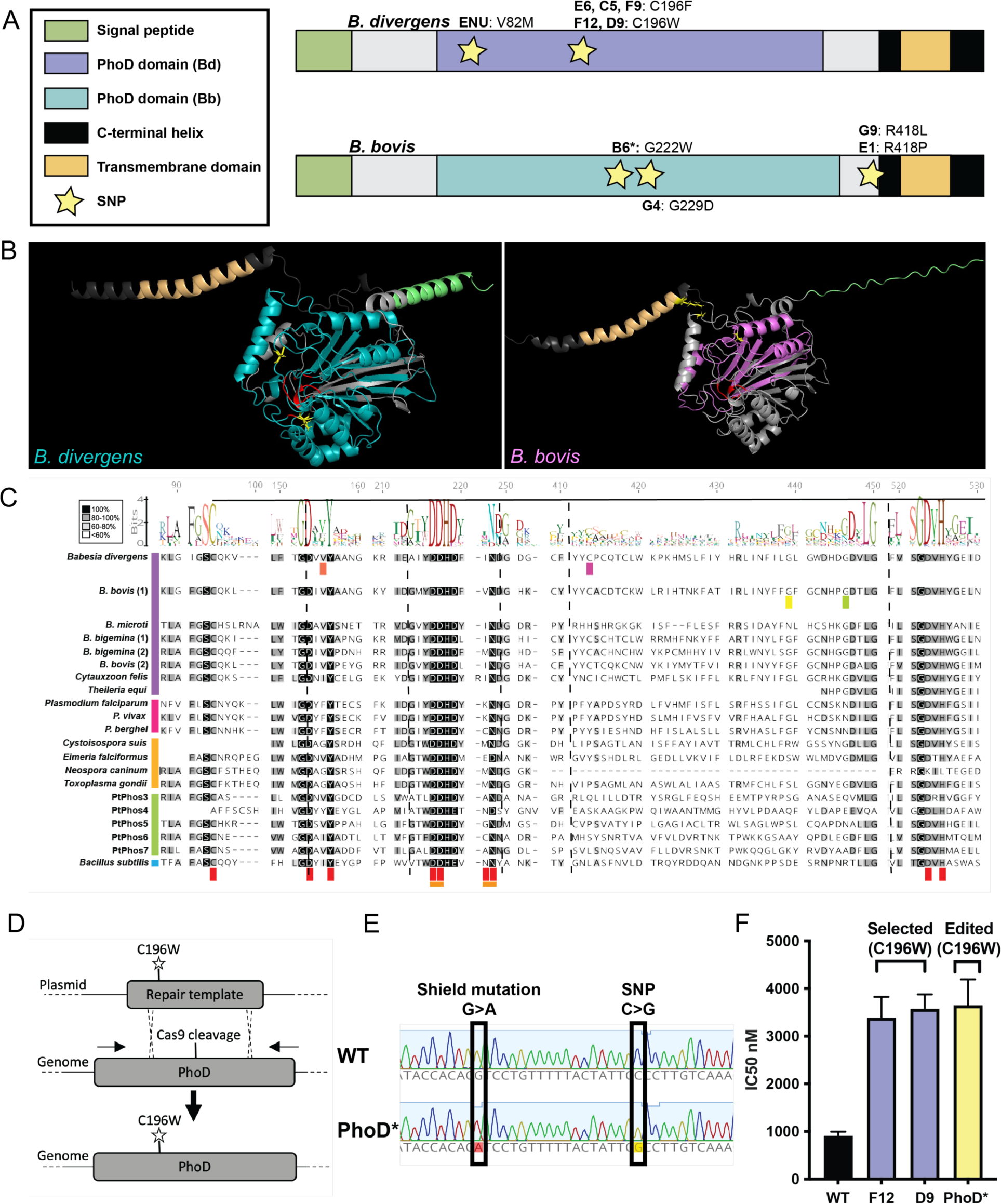
PhoD is a previously uncharacterized, highly conserved alkaline phosphatase which confers resistance to MMV019266. (A) Babesia phoD gene model schematics. mutation locations are denoted by yellow stars. *B. divergens* (Bdiv_001570c, top); *B. bovis* (BBOV_I003300, bottom), key to the left denotes the different predicted domains of the protein. (B) Alphafold structure of Bdiv_001570c (left) and BBOV_I003300 (right). The functional domains are highlighted: signal peptide (green), phoD domain (teal (*B. divergens*), purple (*B. bovis*)), C-terminal helix (dark gray), transmembrane domain (orange). Mutations are highlighted in yellow in each structure and shown in stick form. The active residues of the phosphatase domain are highlighted in red. (C) Alignment of the phoD domain (InterPro). Sections of the protein alignment of the predicted phoD domain which contain mutations, metal binding residues (bottom, red), or the active site (bottom, orange). Each residue where a SNP has occurred is marked by a different color: BdPhoD-V82 (salmon), C196 (magenta); BbPhoD G222 (yellow), G229 (lime green). BbPhoD R418 is not shown as it is outside the phoD domain. Apicomplexans are arranged as follows by order and denoted by colored bar to the right of the species name: piroplasmida (purple), haemosporidia (pink), Eucoccidiorida (orange). PhoDs from *Phaeodactylum tricornutom* are denoted by the green bar (PtPhos3-7). Bacteria (Bacillus subtilis) is denoted by blue. The consensus amino acid residue is denoted along the top of the alignment. Sequence gaps are denoted by vertical hashed lines. The legend for the percent similarity (Blossum45) coloring in the alignment is in the upper left corner. (D) Schematic of the CRISPR/Cas9 strategy used to introduce C196W into B. divergens. (E) Sanger sequencing confirms the presence of the SNP resulting in the C196W mutation, as well as the shield mutation introduced to remove the PAM site to prevent further cutting. (F) Testing of the CRISPR engineered clone (yellow) reveals a shift in IC50 that is the same as those observed in the in vitro selected lines (purple). Error bars represent standard deviation.

In addition to these parasites selected through *in vitro* evolution, we also used ENU mutagenesis on *B. divergens* to assess if mutations would occur in Bdiv_001570c. After exposure to ENU, parasites become highly resistant (∼10 X IC_50_ to MMV019266) after ∼2 weeks post treatment (data not shown). Sequencing of BdPhoD in two resistant clones, one from the 1 mM selection and the other from the 1.5 mM selection, revealed the presence of a novel mutation, V82M, further supporting this gene as the causative agent of resistance.

This gene is annotated as a putative membrane protein with a C-terminal transmembrane helix, containing a phoD-like phosphatase domain (**Fig 2A,B**). This configuration is observed in several phoD phosphatases in the marine diatom *Phaeodactylum tricornutum* [66]. BdPhod shares the highest similarity with the *P. tricornutum* PtPhos6, while BbPhoD is most similar to PtPhos5 and PtPhos7 (**SI Fig 1C**). The gene is homologous to alkaline phosphatase (phoD-like) in both *P. falciparum* and *T. gondii*. Interestingly, this gene also contains a predicted signaling peptide, which is predicted by PATS [87] and ApicoAP [88] to be an apicoplast signal peptide. BbPhoD and BdPhoD are homologous to a protein in *P. falciparum* identified in the apicoplast proteome [89] (**Fig 2 C**). The phosphatase domain is highly conserved within the *Babesia* genus, and also across apicomplexans, marine algae, and bacteria despite differences in overall gene architecture (**SI Fig 1A**). Furthermore, protein sequence alignments reveal high conservation across apicomplexans, diatoms, and bacteria in the metal binding residues and predicted active site, suggesting these phoD phosphatases are functional in apicomplexans (**SI Fig 1B**). Differences in gene architectures of phoD phosphatases is observed in both bacteria and diatoms, and this variation is hypothesized to determine the localization of the protein in the cell [64,65]. In bacteria, phoD phosphatases are monomeric enzymes that catalyze the hydrolysis of both phosphomonoesters and phosphodiesters [63]. In both bacteria and marine algae, phoD is involved in salvaging inorganic phosphate during phosphate starvation [60,63,67,90]. PhoD may serve a similar role in *Babesia,* and the identification of mutations that may alter catalytic activity suggest that modulations in inorganic phosphate levels are important in generating resistance to MMV019266.

### Modeling of PhoD reveals mutations near predicted catalytic domains and supports mechanism of indirection resistance to MMV019266

Using Alphafold [83] we generated a predicted protein structure of PhoD in both *B. bovis* and *B. divergens*, both with and without the predicted signal peptides (**Fig 2B, SI Fig 2A, B**). In all cases, the truncated and full length peptide sequences yielded very similar folded structures, showing that the signal peptide does not affect the predicted folding of the protein (**SI Fig 2E**). Aligning the protein structures of BdPhoD and BbPhoD shows high structural conservation (RMSD = 0.833) (**SI Fig 2C**). The majority of the mutations that were selected for are located proximal to the predicted active site in *B. divergens*, and around the base of the C-terminal helix in *B. bovis* (**SI Fig 2D**). In BdPhoD, V82M is directly next to two of the predicted, conserved metal binding residues (D80, Y83). C196W/F is proximal to the predicted active site of BdPhoD (**SI Fig 2D**). The selection for a mutation of C196 to two different amino acids (W or F) independently, with different levels of resistance, suggests that the replacement of cysteine with phenylalanine results in higher resistance than tryptophan, and that C196F may have a lower catalytic activity than C196W. The location of mutations suggest they have an impact on catalytic activity. Conversely, in BbPhoD all appear to be located closer, or within, the C-terminal helix-suggesting these mutations may alter substrate access. Thus, it is likely that resistance to MMV019266 mediated through these polymorphisms is directly linked to the level of activity of phoD.

PhoD is a unique alkaline phosphatase in that it’s catalytic activity relies on Fe^3+^ and two Ca^2+^ ions, rather than zinc ions in canonical alkaline phosphatases [60–64]. Access to the active site of phoD is controlled by a C-terminal helix [62]. Unfortunately, the crystal structure of phoD was resolved in the closed conformation, and despite efforts the topology of the open structure or the mechanism of movement of the C-terminal helix has not been resolved. Despite this, we attempted molecular docking to assess the ability of MMV019266 to interact with BbPhoD and BdPhoD. We did not identify any high confidence docking sites in either species or in *B. subtilis*. Together, these data indicate that there is no direct interaction of MMV019266 with the active site of phoD, supporting the role of the protein as a resistance mechanism.

### Reverse genetics validates *phoD* mutations in resistance to MMV019266

To confirm that selected mutations in BdPhoD confer resistance to MMV019266, we adapted the previously successful CRISPR/Cas9 system of gene editing to *B. divergens* to introduce the single point mutation C196W into wild type parasites (**Fig 2 D, E**) [57]. Position 196 in BdPhoD was mutated in two independent selections with two different amino acid substitutions (C196W, C196F). C196W was selected for analysis as it is both the larger of the two amino acid substitutions, and appeared only in one independent selection to eliminate the possibility that C196F was a hitchhiking mutation. Clonal parasite lines containing the single point mutation were resistant to MMV019266 at a level consistent with that observed from clones generated via intermittent selection, showing this mutation is alone sufficient for resistance (**Fig 2F**). Upon validation, we hereafter refer to Bdiv_001570c and BBOV_I003300 as BdPhoD and BbPhoD, respectively.

### Perturbing the abundance of BdPhoD alters the level of resistance

We generated a line of parasites which episomally overexpress BdPhoD on its endogenous promoter (**Fig 3A**). Overexpression of BdPhoD did not induce a proliferation defect (**Fig 3B**). Five fold over expression of BdPhoD was demonstrated by qRT-PCR (**Fig 3C**). Protein expression of phoD-HA in the overexpression line was confirmed by immunoblotting (SI Fig 5). Dose-response assays to MMV019266 we performed on overexpression parasites in comparison to wild type parasites (**Fig 3D**). We observed a 3-fold decrease in IC_50_ in the parasites overexpressing BdPhoD, revealing that an increase in the protein level leads to increased sensitivity to MMV019266. While direct targets may have complex mechanisms of action resulting in similar phenotypes observed here (i.e. DNA gyrase and ciprofloxacin [91]), these results suggest that BdPhoD is likely a resistance mechanism, rather than the direct target of MMV019266.

**Figure 3.**
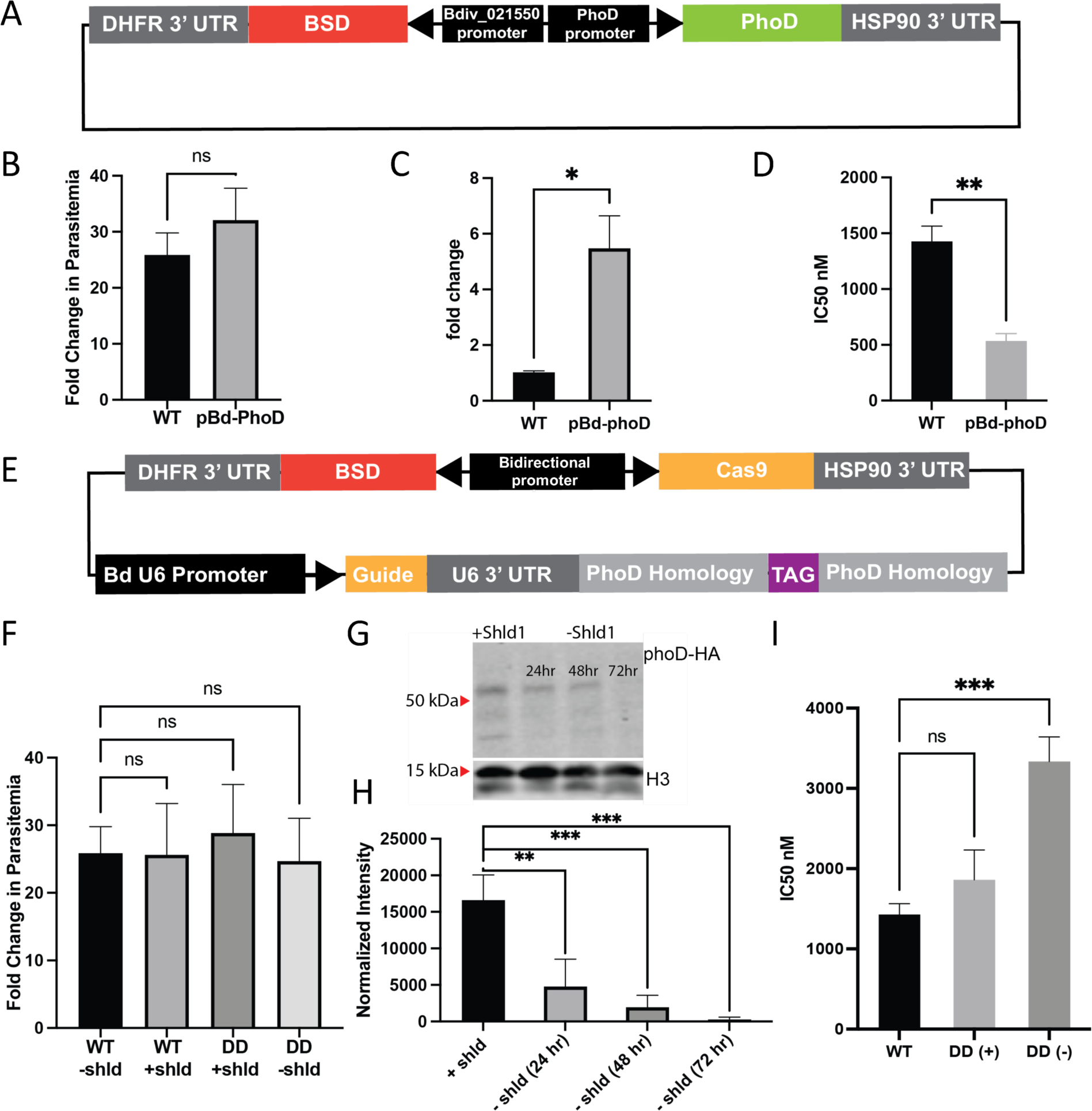
Perturbations to the level of BdPhoD lead to changes in resistance level. For significance values: NS p > 0.05, * p < 0.05, ** p < 0.005, *** p < 0.0005. All error bars represent standard deviation. (A) Schematic of the plasmid used to overexpress PhoD-HA. (B) Proliferation of wild type (WT) and overexpressing parasites (pBd-PhoD) measured in fold change in parasitemia after 48 hours of growth, counted on thin smears by light microscopy (n = 3). Significance determined by paired t-test. (C) qPCR of phoD in WT and Bd-phoD parasites (n = 3). Significance determined by paired t-test. (D) IC_50_ for MMV019266 in WT and Bd-phoD parasites. (E) Schematic of the plasmid used to tag BdPhoD with an HA-DD-glmS tag for inducible knockdown. (F) Proliferation of wild type (WT) and overexpressing parasites (pBd-PhoD) measured in fold change in parasitemia after 48 hours of growth, counted on thin smears by light microscopy (n = 3). Significance determined by one way ANOVA. (G) Immunoblot for ⍺-HA to detect PhoD-HA. Knockdown (-Shld) was followed for 72 hours. Loading control is ⍺-Histone H3. (H) Quantification of the immunoblots following knockdown of BdPhoD (-Shld), normalized to loading control. Significance determined by one way ANOVA. (I) IC_50_ for MMV019266 in WT and DD (+Shld) and (-Shld) parasites. Significance determined by one way ANOVA.

To further assess the interaction between MMV019266 and BdPhoD, we generated an inducible knockdown parasite line by tagging the gene with a destabilization domain (DD), which has been used with success previously in *B. divergens* [57] (**Fig 3E, SI Fig 5**). In this system, the protein is stabilized by supplementation of the culture media with Shld1, and knock down is induced upon washout via protein degradation [57]. Assessing parasite proliferation over 72 hours showed no significant difference between the knock-down and wild type parasites, suggesting the level of knock-down of BdPhoD achieved does not confer a significant phenotype on asexual blood stage parasites (**Fig 3F**). After removing Shld1, a 3.5 fold reduction in protein is observed via immunoblot by 24 hrs post wash out, and the protein reached 62.5 fold reduction by 72 hrs post Shld1 wash out (**Fig 3G**). Dose-response assays to MMV019266 were performed on Shld1 (+) and Shld1 (-) parasites, in comparison to wild type parasites. We observed a 2.3 fold increase in IC_50_ in the Shld1 (-) knock-down parasites, showing that depletion of the protein results in resistance to MMV019266 (**Fig 3H**). There was a non-significant increase in IC_50_ in the Shld1 (+) condition, which suggests slight knockdown due to the presence of the destabilization domain tag. Taken together, these results demonstrate that modulating the protein levels of BdPhoD alters resistance to MMV019266, consistent with BdPhoD acting as a resistance mechanism.

Finally, we endeavored to generate a knockout. Very few essential genes have been validated in *Babesia,* and to date, no large screen of essentiality has been done. As such, essentiality of BdPhoD has yet to be described. In *P. falciparum*, the homologous gene (PF3D7_0912400) did not display a growth defect in the *piggybac* transposon based screen [92]. No data is reported on essentiality in *P. bergei* in the plasmoGEM screen [93]. In a genome-wide CRISPR screen in *T. gondii* the homologous gene (TGME49_265830) has a modestly negative phenotype score, indicating a minor growth defect upon disruption, suggesting this gene is nonessential in *T. gondii* [94]. In this study, we have shown there is no significant defect on growth rate or morphology upon knockdown of BdPhoD (**Fig 3F**). We were unable to generate a knockout of BdPhoD despite multiple efforts. Amplification of the BdPhoD locus (primers designed outside of the homology regions used) in clonal lines from knock-out transfections revealed that parasites always retained the wild-type BdPhoD locus, in addition to integrating the knockout cassette, suggesting a genome arrangement occurring to prevent knock out of the gene (**SI figure 3**). We also observed no loss of function mutations in phoD in either species of our selected lines (**SI table 4**). These data suggest PhoD may perform an essential function in *B. divergens* which can be maintained at very low levels of expression, but cannot be entirely ablated in the parasite. Interestingly, knock out of phoD in the diatom *P. tricornutum* results in a significant growth phenotype, suggesting a more essential role for the gene may be an ancestral trait [67].

### BdPhoD is localized in the endoplasmic reticulum and apicoplast

In both prokaryotes and eukaryotes, phoD localization varies widely depending on gene architecture and functional requirements of the cell [64–66]. The diversity of locations of phoD between suggest divergent function based on the phosphorous needs of the particular species. In diatoms, several phoD-like phosphatases exist: of these, PtPhos5 and PtPhos6 share the highest sequence similarity with BdPhoD and BbPhoD, respectively (**Fig 4L, SI Fig 1C**). Consistent with the sequence predictions in *Babesia,* both PtPhos5 and PtPhos6 are predicted to be a membrane integrated phosphatase with a single transmembrane helix. PtPhos5 is localized to the plastid and endomembrane system, while PtPhos6 was detected in the ER [66]. In apicomplexans, phoD was identified in the apicoplast proteome of *P.* falciparum, and in the endoplasmic reticulum proteome fraction of *T. gondii* [89,95,96]. As PhoD in *Babesia* has a predicted apicoplast signal peptide (as predicted by PATS [87]), we predicted we would identify localization in the apicoplast, and based on homology, with the endoplasmic reticulum. Using the CRISPR/Cas9 system described above, we introduced a GFP tag to the 3’ end of *phoD* at the endogenous locus in *B. divergens* (Bd-phoD-GFP) (**Fig 4A, SI Fig 5**). Through live cell imaging, we found that phoD-GFP was compartmentalized, adjacent to the nucleus (**Fig 4G**). To determine which organelle(s) contained BdPhoD, Bd-phoD-GFP was analyzed by immunofluorescence and live cell imaging with colocalization markers. To provide a level of quantification the amount of overlap between signals was assessed using a qualitative metric for high, mid, and low level of overlap between the fluorescent signals for at least 20 1N and 2N parasites, which is described in **Fig 4B**.

**Figure 4.**
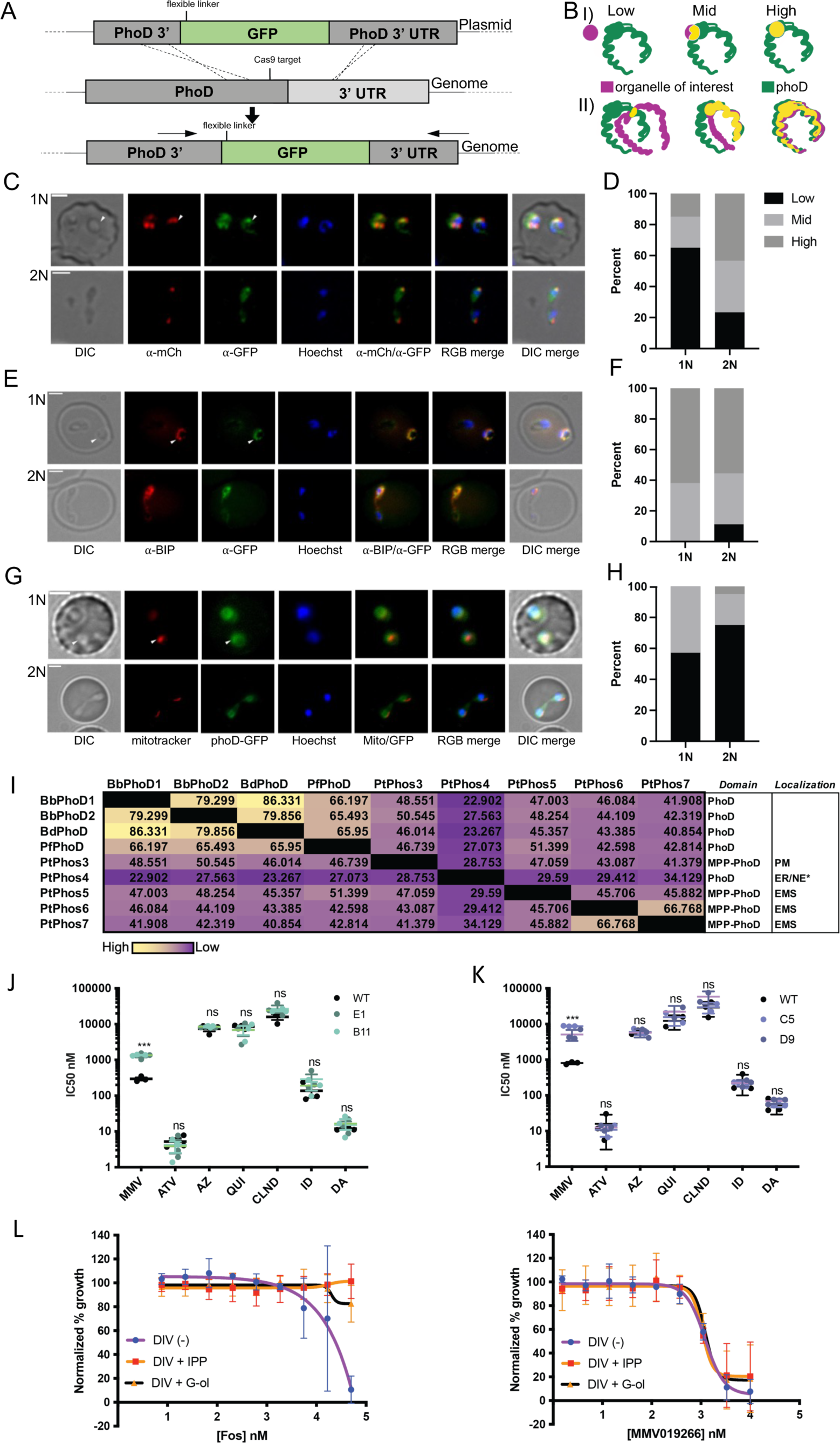
PhoD is compartmentalized to the ER and apicoplast in the cell, is dynamic depending on cell state. For significance values: NS p > 0.05, * p < 0.05, ** p < 0.005, *** p < 0.0005. (A) A schematic of the endogenous tagging strategy used to introduce the GFP tag for localization. Arrows indicate location of primers used for validation of tag integration. (B) A schematic of the grading used to quantify the levels of colocalization between various organelles. Magenta represents the organelle of interest, green represents phoD. I) shows the grading scheme for colocalization with the mitochondrion and apicoplast. II) shows the grading for colocalization with the ER. (C) Fixed and permeabilized transgenic cells (endogenous tagged phoD-GFP) expressing ectopic lytB-mCherry (pBd-SP+TP_lytB_-mCherry) stained using ⍺-GFP (phoD) and ⍺-mCherry (red, lytB – apicoplast marker). Top row of images are 2N parasites, bottom row are 1N parasites (white arrow indicates the 1N parasite of interest). Scale bar represents 2 µm. (D) Quantification of C, number of cells counted and graded – 1N = 20, 2N = 30. (E) Fixed and permeabilized transgenic cells (endogenous tagged phoD-GFP) stained using ⍺-GFP (green, phoD) and ⍺-BIP (red, endoplasmic reticulum marker). Top row of images are 2N parasites, bottom row are 1N parasites (white arrow indicates the 1N parasite of interest). Scale bar represents 2 µm. (F) Quantification of E, number of cells counted – 1N = 21, 2N = 27. (G) Fixed and permeabilized transgenic cells (endogenous tagged phoD-GFP) stained using ⍺-GFP (green, phoD) and ⍺-BIP (red, endoplasmic reticulum marker). Top row of images are 2N parasites, bottom row are 1N parasites (white arrow indicates the 1N parasite of interest). Scale bar represents 2 µm. (H) Quantification of G, number of cells counted – 1N = 21, 2N = 20. (I) Cross resistance testing for two clones of resistant to MMV019266 B. bovis parasites (selection A-B11; selection B-E1) Compound legend is as follows: MMV2-MMV019266, MMV3-MMV085203, ATV-atovaquone, AZ-azithromycin, QUI-quinine, CLIND-clindamycin, ID-imidocarb dipropionate, DA-diminazene aceturate. Black dots represent WT IC_50_, colored dots represent resistant clone IC50. Statistical significance calculated by ordinary one-way ANOVA for each compound. Error bars represent SD. (J) Cross resistance screening in B. divergens. Formatting is as in panel I. Statistical significance calculated by ordinary one-way ANOVA for each compound. (K) Dose-response curve of fosmidomycin (right, positive control) and MMV019266 (left) in B. divergens using either geranylgeraniol (G-ol, black line) or IPP (orange line). N = 3 biological replicates. Error bars represent standard deviation between 3 independent dose-response curves. (L) Similarity matrix of apicomplexan and *Phaeodactylum tricornutum* phoD domains based on protein alignments (similarity by % similarity Blossum45, threshold 0): BbPhoD1 = BBOV_I003300, BbPhoD2 = BBOV_I003305, BdPhoD = Bdiv_001570c, PfPhoD = PF3D7_0912400. The five phoD genes from *P. tricornutum* are included (SI Table 2). The type of InterPro predicted phosphatase domains are included in the first right column, localization in the last right column. Localization abbreviations: PM= plasma membrane, ER= endoplasmic reticulum, NE= nuclear envelope, EMS= endomembrane system (membrane surrounding ER, plastid).

We first investigated the potential for apicoplast localization. Previously, LytB was shown to localize to the apicoplast in *B. bovis,* and we designed similar constructs to assess this tool for localization of the apicoplast in *B. divergens* [71]. As previously done in *B. bovis*, we fused the signal peptide and transit peptide of LytB from *B. divergens* to GFP and observed its localization via live cell imaging (**SI Fig 6B**). The localization of SP+TP_lytB_-GFP was consistent with what was observed previously in *B. bovis* [71], and appeared as a punctate structure that did not overlap with the mitochondrion as expected, demonstrating specific localization to the apicoplast (**SI Fig 6B**). Due to the markerless nature of the CRISPR/Cas9 gene editing, we were then able to co-transfect our endogenously tagged line phoD-GFP with the episomal marker of the apicoplast (SP+TP_lytB_-GFP). In order to do this cotransfection, we swapped GFP with mCherry in the original pBdEF-SP+TP_lytB_-GFP-BSD with SP+TP_lytB_-mCherry to assess colocalization of PhoD with the apicoplast. We also confirmed that mCherry localization was the same as GFP (**SI Fig 6C**). There is an overlap of signal between the apicoplast and PhoD, particularly in the punctate signal from BdPhoD. This suggests there is colocalization with the apicoplast. However, there is BdPhoD signal outside of the apicoplast as well, suggesting a localization and function beyond the apicoplast (**Fig 4C**). Furthermore, the level of overlap with the apicoplast is higher in 2N parasites than in 1N parasites, suggesting the localization of BdPhoD is dynamic in the cell (**Fig 4D**).

With the knowledge that BdPhoD localizes to the endomembrane system *in P. tricornutum* we next investigated colocalization with the ER [66]. The signal beyond the apicoplast showed a horseshoe shape that was excluded from the nucleus, typical of the endoplasmic reticulum. Based on the reports of phoD being located in the ER in *T. gondii* [95] and marine algae [66,97] we next used the phoD-GFP line to perform colocalization with the ER specific marker BIP. PhoD-GFP overlaps with the ER specific marker, indicating colocalization. Interestingly, the localization of BdPhoD was dynamic, and changed from a diffuse horseshoe with a distinct punctate shape in the 2N (newly divided), to a more diffuse and elongated distribution between in the 1N (**Fig 4E**). The level of overlap between BdPhoD and the ER in 1N and 2N parasite is similar (**Fig 4E,F**). Of note, apicoplast proteins are also trafficked through the endoplasmic reticulum [98–100], so this localization may be indicative of more active protein trafficking immediately post-division. A similar pattern of fluctuation is observed with the BIP ER marker, further suggesting a connection between the apicoplast and ER in *B. divergens*. Interestingly, the ER and apicoplast have been shown to be in direct contact in *T. gondii* [101], there may be a direct flow of nutrients, metabolites, or messengers such as inorganic phosphate between the organelles. Together, these results suggest that BdPhoD is localized to the apicoplast and endoplasmic reticulum-this may be a pattern of trafficking or differential localization based on cellular state, and may indicate BdPhoD has functions in multiple areas in the cell.

Finally, we wanted to assess possible co-localization with the mitochondrion. Previous reports have observed some apicoplast proteins containing signal peptides that dually target the protein to both the plastid and the mitochondrion [102–105]. Using the mitochondrial stain MitoTracker Red CM-H2XRos we show that BdPhoD does not colocalize with the mitochondrion (**Fig 4 G,H**). To ensure the endogenous GFP tag was not disrupting localization, we used the predicted signal peptide and transit peptide of phoD fused to GFP for comparison in combination with mitotracker. Like the endogenously tagged line (Bd-phoD-GFP), episomal SP+TP_phoD_-GFP has horseshoe like localization that does not overlap with the mitochondrion (**SI Fig 6A**).

Together, we have demonstated that BdPhoD localizes to the apicoplast and ER in *B. divergens.* Aligning both the full gene and the phoD domain of BdPhoD and BbPhod to the 8 alkaline phosphatases (5 phoD, 2 phoA, 1 phytase) reveals that the phoDs with localization to the endomembrane system share the highest protein sequence similarity (**Fig 4I, SI Fig 1C**). Whether the catalytic domain of BdPhoD is exposed to the cytoplasm or to the ER lumen is unknown. This localization pattern was observed in PtPhos5 *P. tricornutum* leading us to hypothesize that BdPhoD functions in a similar role [66]. This suggests that *Babesia* PhoD has been retained for functions related to the endoplasmic reticulum and apicoplast, and likely is involved in mobilization of intracellular inorganic phosphate.

### Protein alignment of PhoD across apicomplexa reveals reduction in diversity

Searching the plasmoDB, piroplasmaDB, and toxoDB databases we identified a total of 128 phoD domain containing genes across 53 species. These results contain multiple strains of certain species (e.g. *T. gondii, P. falciparum*). Of these, 19 genes were identified in 12 *Babesiidae* and *Theileriidae* species, 56 genes in 22 species of *Plasmodiidae* spp., and 53 genes in 12 species of *Eimeriidae* and *Sarcocystidae* (**SI Table 2**). In the piroplasms, organisms either contained one or two genes with an annotated phoD domain. The majority of *Plasmodium* species have a single gene containing a phoD domain. Finally, in *Eimeriidae* and *Sarcocystidae* the majority of organisms have two genes containing a phoD domain, with the exceptions being *Besnoitia besnoiti* having three. No orthologs were identified through BLAST of phoD and phytase from *P. tricornutum*. *P. tricornutum* is annotated to contain 5 phoD like alkaline phosphatases, each with different localizations. In all of the phoD domain containing genes identified, none have evidence of being secreted. This suggest phoD was retained in apicomplexans to serve functions within the cell, and that the need for secreted phoD or phoD facing the external environment in the plasma membrane has been lost.

### Treatment with MMV019266 does not result in a delayed death phenotype and cannot be rescued with isopentenyl pyrophosphate (IPP) or geranylgeraniol (G-ol)

Given the apicoplast localization of BdPhoD, we next investigated the possibility of a delayed death phenotype with MMV019266-something characteristic of apicoplast targeting compounds [106,107]. We measured the nuclear content of synchronized parasites treated with MMV019266 (10 X IC_50_ = approx. IC_90_) over the course of 12 hours as a proxy for understanding killing speed [108]. Twelve hours was selected as a representation of a single intraerythrocytic development cycle [108]. The synchronization protocols for *Babesia* are imprecise and result in the majority of parasites starting at 1N, but a remained of parasites which failed to be mechanically released from the cell and will start the time course later in the cell cycle (2N), which can be observed by nuclear content-approximately 20% of the total parasites at the 0 hours post invasion (hpi) are still at 2N, rather than having been efficiently released and reinvaded. Regardless, over the course of 12 hours in both *B. divergens* and *B. bovis* we observed no increase in the number of 2N parasites, and a decrease in the number 1N parasites which correlates with an over all decrease in parasitemia (**SI figure 7A, B**). Thus, MMV019266 prevents parasites from dividing and causes rapid death. Together, the phenotypic data suggests MMV019266 kills *B. bovis* and *B. divergens* within the first replication cycle.

To assess the specificity of the resistance generated by *in vitro* evolution, we tested resistant parasites against a panel of clinically relevant compounds (diminazene aceturate, imidocarb dipropionate, clindamycin, quinine, azithromycin, atovaquone) to assess for potential cross resistance. Of note, clindamycin targets the apicoplast [109]. We observed no cross resistance to any compounded tested supporting a specific mechanism of resistance generated against MMV019266 (**Fig 4 J,K**).

Finally, we attempted to rescue MMV019266 treatment through supplementation of media with isopentenyl pyrophosphate (IPP) and Geranylgeraniol (G-ol). Rescue of compounds using IPP is a well-established method for confirming apicoplast targeting [107]. Inhibition of the apicoplast thus should be rescued by supplementation of the IPP or G-ol; rescue has been demonstrated in *B. orientalis* and *B. microti* [110,111]. Using the apicoplast targeting compound fosmidomycin (inhibitor of isoprenoid biosynthesis) as a control, we were able to confirm the ability to rescue with both geranylgeraniol and IPP in *B. divergens.* Treatment with MMV019266 could not be rescued in *B. divergens* (**Fig 4L**). These results further support phoD as a resistance mechanism of MMV019266, rather than the direct target, and that the direct target is not in the apicoplast.

## Discussion

Current treatments for babesiosis in both humans and animals are suboptimal and the efficacy of the same compound can vary significantly across species, hindering treatment efforts. In this study we have developed a comparative chemical genomics pipeline, demonstrating the power of using multiple species in *in vitro* evolution experiments to identify conserved genes involved in resistance. We identified a compound of interest MMV019266: a thienopyrimidine compound with demonstrated anti-malarial activity with no validated target [112]. In mycobacteria, thienopyrimidines have been shown to be activated by nitroreductases, releasing damaging nitric oxide [113]. However, thienopyrimidine scaffolds have been shown to target a variety of enzymes across prokaryotes and eukaryotes, and thus no single conserved mechanism is known [114]. Previously, MMV019266 was identified to block transmission in *P. falciparum* [115], and has been demonstrated to have specific activity against *P. falciparum* gametocytes *in vitro* [116]. Based on this knowledge, MMV019266 was an attractive compound candidate for *in vitro* evolution in *Babesia.* Through CCG, we identified and validated phoD, an alkaline phosphatase, as a resistance determinant in *Babesia*.

At the outset of this study, no known target of MMV019266 had been identified: recently however, a evidence of a potential target was described in *P. falciparum* using in vitro evolution. Mutations in cytoplasmic isoleucyl tRNA synthetase (PfcIRS) were identified after selections using other thienopyrimides which were shown to confer resistance, including to MMV019266 [48]. Further, knockdown of PfcIRS was shown to sensitize the parasites to these compounds. We did not identify mutations in the *B. divergens* or *B. bovis* ortholog of PfcIRS, nor were phoD mutations identified in the *P. falciparum* selections. Mutations in phoD were the only ones conserved in both *Babesia* species. As such, we cannot confirm that cIRS is the target in *Babesia,* however this finding in *P. falciparum* further supports our conclusion that mutations in phoD serve as a resistance mechanism, rather than as the direct target of the compound.

PhoD is a previously uncharacterized alkaline phosphatase (AP) in *Babesia*. PhoD family phosphatases have been extensively characterized in *Bacilllus subtilis*, as well as in the cyanobacteria *Aphanothece halophytica* and *Synechococcus elongatus* [60,62,63]. The family of phoD alkaline phosphatases has been shown to be the most abundant of the alkaline phosphatases in marine bacteria [64]. In these prokaryotes phoD genes are predicted to localize to different subcellular compartments including the periplasm, cell membrane, cell wall, and secreted into the extracellular space depending on the species and sequence architecture [60,61,63,64]. In eukaryotic phytoplankton such as dinoflagellates and diatoms, APs have also been shown to have highly variable localizations, and demonstrates rapid evolution which may be driven by different strategies for accessing dissolved organic phosphate [65,66]. The diatom *P. tricornutum* has three families of AP which perform important cellular functions: phoD, phoA, and phytases [66]. Of these, only phoD is conserved in apicomplexans. In both marine prokaryotes and eukaryotes, there are secreted phoD phosphatases [64–66]. We found no evidence for secretion in *B. divergens,* and it appears that these secreted phoD phosphatases have been lost in apicomplexa, and that there has been generally a loss in phoD diversity in the parasites-a possible adaptation to a parasitic lifestyle.

Alkaline phosphatases (AP) play important roles in phosphate starvation [64,68]. In *B. subtilis,* phoD expression is suppressed during vegetative growth, and is upregulated during phosphate starvation [60]. PhoD is similarly upregulated during phosphate starvation in *A. halophytica* and *S. elongatus* [63]. Further, in the diatom *P. tricornutum,* several APs are upregulated during phosphate starvation [66]. While much remains unknown about responses to phosphate starvation in apicomplexan parasites, nearly all key metabolic processes depend on the availability of both organic and inorganic phosphorous. For example, phosphate translocators are essential to apicoplast function in *T. gondii* [117,118]. Further, phosphate is essential for metabolic processes occurring in the ER [101,119,120]. Recently, phosphate starvation in *Toxoplasma gondii* was shown to restrict growth. Yet, this restrictive environment did not lead to the upregulation of phosphate transporters, suggesting that parasites are able to access internal stores of phosphate, such as phospholipids on the plasma membrane, by an unknown mechanism [121]. Membrane bound phoD has been shown to serve a very similar role in bacteria-interacting with the cell wall to release inorganic phosphate during times of phosphate starvation [60,122]. PhoD may act in a similar fashion during phosphate starvation in *Babesia*, acting to mobilize inorganic phosphate for processes such as isoprenoid biosynthesis, ATP biosynthesis, and membrane transport. Decreasing the abundance of phoD may lead to a decrease in phosphate scavenging activity within the cell, resulting in less free inorganic phosphate for metabolic activities.

The diatoms *P. tricornutum* and *Thalassiosira pseudonana* are known to replace phospholipids with non-phospholipids during times of phosphate starvation. As the endomembrane localized PtPhos6 in *P. tricornutum* is strongly upregulated under phosphate starvation conditions, it has been hypothesized that these endomembrane localized phoD phosphatases may function in degredation of phospholipids in phosphate limited conditions[66]. Altering the balance between phosphorylated and non-phosphorylated lipids in a membrane can alter the permeability of the membrane. We speculate that the catalytic domain of phoD in *Babesia* serves to dephosphorylate the phospholipids in the membranes surrounds the ER and apicoplast. While we do not know the orientation of the catalytic domain, we hypothesize that if the target is in the cytoplasm (such as cIRS[48]), the catalytic domain points towards the cytoplasm. Conversely, if the target is in the ER or apicoplast, the catalytic domain would point into the lumen. In doing so, the enzyme effectively alters the permeability of these membranes, sequestering the drug away from the target (either the cytoplasm or ER/apicoplast lumen) (**SI Fig 8**). Our data suggest that decreasing the catalytic activity of phoD leads to increased resistance. In this model, decreased catalytic activity would lead to a decrease in dephosphorylation of phospholipids in the membranes, altering the membrane permeability (**SI Fig 8**). If the drug is sequestered in the ER, the drug may be detoxified by ER resident enzymes, or effluxed from the parasite via vesicular transport. Alternatively, decreasing catalytic activity of phoD may alter the levels of free inorganic phosphate in the cytoplasm, leading to broad changes in cellular metabolism which may confer resistance. PhoD appears to broadly impact in general cellular metabolism in *P. tricornutum*. Recently, phoD (PhoD_45757) was shown to play an important role in global cellular function and cell cycle progression [67,97]. Through the generation of a functional knock out, phoD was determined to regulate phosphate uptake and storage, dampen excessive carbon fixation, and restrain nitrogen assimilation in *P. tricornutum* [67]. If phoD plays a similar role in *B. divergens,* it is possible that MMV019266 interferes with a key metabolic process (such as phosphate cycling), and that resistance mutations are able to overcome these disruptions by modulating the levels of free inorganic phosphate in the cytosol. Studying the global metabolic changes in the cell after phosphate starvation or MMV019266 treatment at the transcriptomic, proteomic, and metabolomic levels would address if phoD has involvement in phosphate metabolism related to these environmental perturbations.

In summary, we have presented a MMV019266 as an effective pan-babesiacidal compound. We have identified and validated phoD, a highly conserved alkaline phosphatase, as a resistance mechanism against MMV019266 in *Babesia*. By leveraging the power of *in vitro* evolution and chemical genomics performed in multiple species in parallel, we have built and validated a robust comparative chemical genomics pipeline for high confidence identification of resistance loci in *Babesia* which can be used for future drug target validation and discovery.

## Acknowledgments

The authors would like to thank the entire Duraisingh group for support and helpful discussions throughout this project. We would like to thank Dr. Ralph Mazitschek for his chemical insights related to this project. Thank you to Dr. M.J. Gubbels and Dr. Klemens Engelberg for guidance on IFA protocols. This work was supported by grant 1R21AI153945 (MD) from the National Institutes of Health. CDK was supported by an AHA pre-doctoral fellowship (#19PRE34380106). The funders had no role in study design, data collection and interpretation, or the decision to submit the work for publication.

