## Supplemental Figures for "Comparative chemical genomics in *Babesia* species identifies the alkaline phosphatase phoD as a novel determinant of resistance"

### SUPPLEMENTARY FIGURES

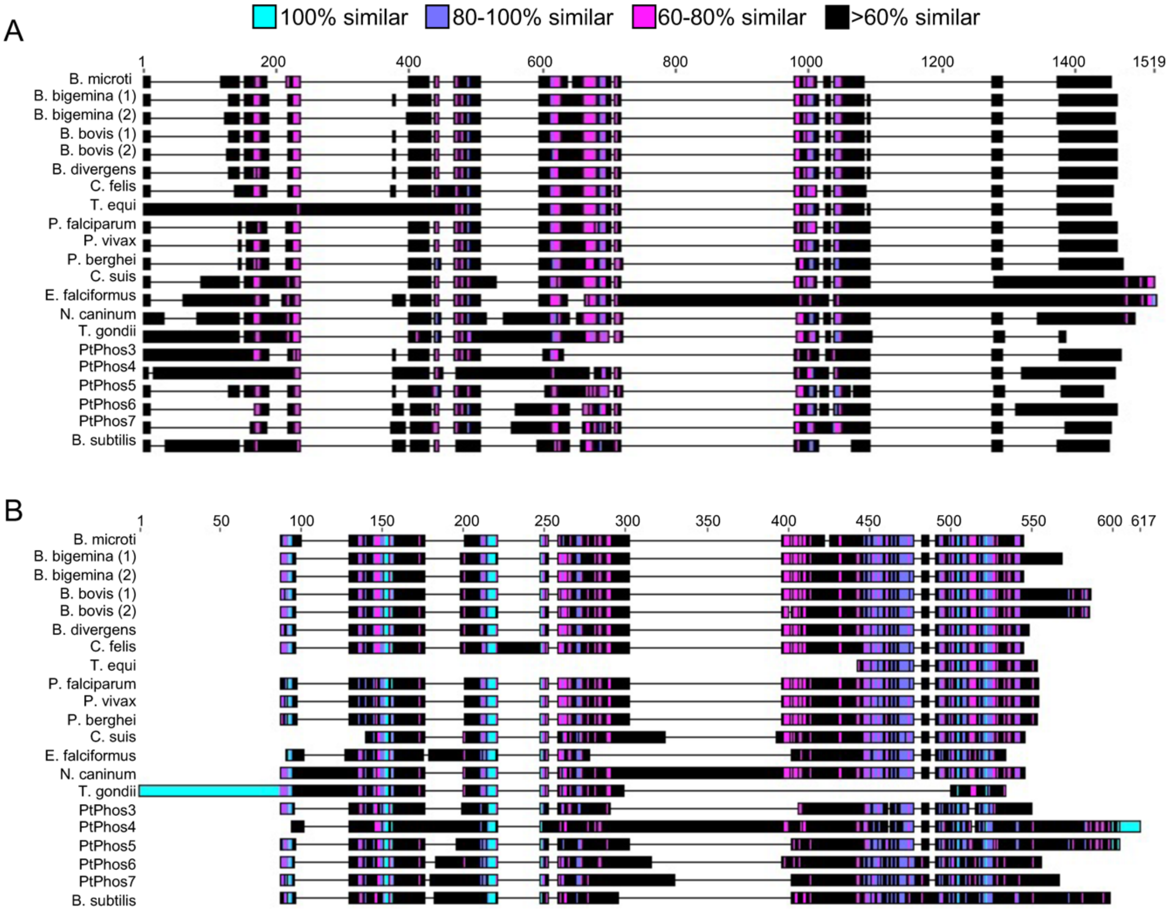


C

| | BbPhoD1 | BbPhoD2 | BdPhoD | PtPhos1 | PtPhos2 | PtPhos3 | PtPhos4 | PtPhos5 | PtPhos6 | PtPhos7 | PtPhos8 | |
|---|---|---|---|---|---|---|---|---|---|---|---|---|
| **BbPhoD1** | | 74.074 | 81.36 | 22.003 | 22.409 | 25.631 | 23.176 | 41.485 | 37.699 | 37.598 | 22.571 | |
| **BbPhoD2** | 74.074 | | 70.806 | 23.182 | 22.689 | 28.35 | 23.647 | 44.348 | 37.148 | 35.225 | 22.048 | |
| **BdPhoD** | 81.36 | 70.806 | | 22.762 | 22.549 | 28.155 | 25.322 | 43.231 | 35.398 | 37.402 | 20.571 | |
| **PtPhos1** | 22.003 | 23.182 | 22.762 | | 31.258 | 30.198 | 35.917 | 22.749 | 27.753 | 31.209 | 33.65 | S |
| **PtPhos2** | 22.409 | 22.689 | 22.549 | 31.258 | | 26.97 | 37.875 | 21.366 | 26.223 | 26.537 | 36.828 | S |
| **PtPhos3** | 25.631 | 28.35 | 28.155 | 30.198 | 26.97 | | 28.241 | 29.787 | 34.924 | 33.397 | 27.744 | PM |
| **PtPhos4** | 23.176 | 23.647 | 25.322 | 35.917 | 37.875 | 28.241 | | 22.173 | 26.667 | 27.343 | 35.349 | ER/NE* |
| **PtPhos5** | 41.485 | 44.348 | 43.231 | 22.749 | 21.366 | 29.787 | 22.173 | | 41.366 | 40.784 | 21.842 | EMS |
| **PtPhos6** | 37.699 | 37.148 | 35.398 | 27.753 | 26.223 | 34.924 | 26.667 | 41.366 | | 63.265 | 26.565 | EMS |
| **PtPhos7** | 37.598 | 35.225 | 37.402 | 31.209 | 26.537 | 33.397 | 27.343 | 40.784 | 63.265 | | 27.231 | EMS |
| **PtPhos8** | 22.571 | 22.048 | 20.571 | 33.65 | 36.828 | 27.744 | 35.349 | 21.842 | 26.565 | 27.231 | | PM |

High — Low

**SI Figure 1. PhoD has high structural diversity across organisms, but has highly conserved residues in the catalytic domain.** A) Full gene alignment of PhoD from various apicomplexa, *P. tricornutum*, and *B. subtilis*. % Similarity is colored calculated by Blossum62: cyan = 100%, purple = 80 – 100%, magenta = 60 – 80%, black = less than 60%. B) Alignment of the InterPro predicted phoD domains, colored by similarity as in panel A. C) Similarity matrix of apicomplexan and *P. tricornutum* alkaline phosphatases (full protein sequence) based on protein alignments (similarity by % similarity Blossum45, threshold 0): BbPhoD1 = BBOV_I003300, BbPhoD2 = BBOV_I003305, BdPhoD = Bdiv_001570c. Yellow represents highest similarity, purple is lowest.

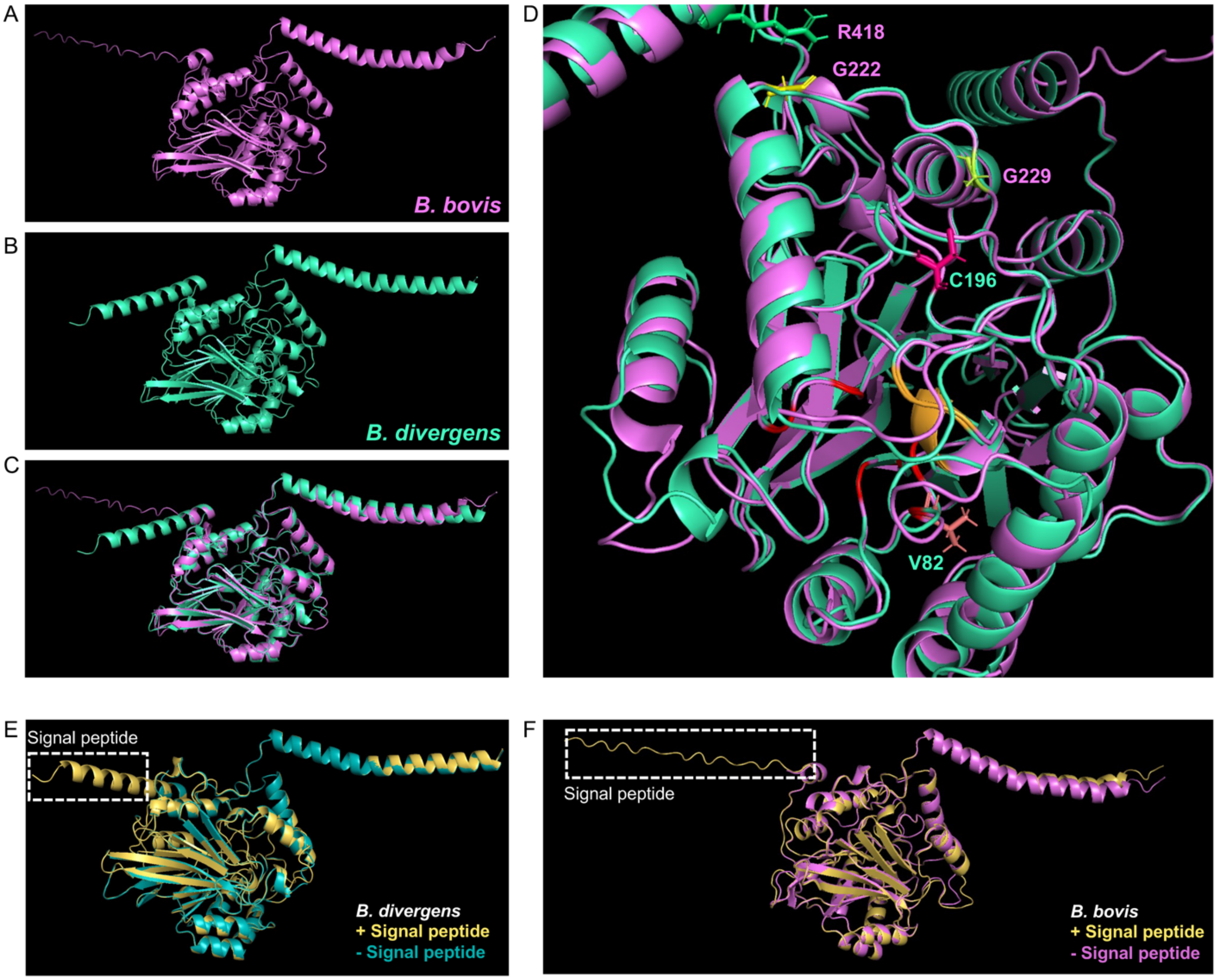


**SI Figure 2.** AlphaFold structures show high structural conservation between species and highlight potential catalytic impact of mutations. (A) AlphaFold structure of full peptide sequence for BBOV_I003300. (B) AlphaFold structure of full peptide sequence for Bdiv_001570c. (C) Alignment of BBOV_I003300 and Bdiv_001570c predicted structures, RMSD = 0.833 (RMSD of 0 is perfect alignment). (D) Zoom of the active site residues of the aligned strucutres, with the SNP mutations highlighted as sticks. *B. bovis* mutations are labeled in purple*, B. divergens* are labeled in teal. (E) Alignment of the AlphaFold predicted structures of Bdiv_001570c with (yellow) and without (teal) the predicted signal peptide (boxed in white). RMSD = 0.190. (F) Alignment of the AlphaFold predicted structures of BBOV_I003300 with (yellow) and without (purple) the predicted signal peptide (boxed in white). RMSD = 0.286.

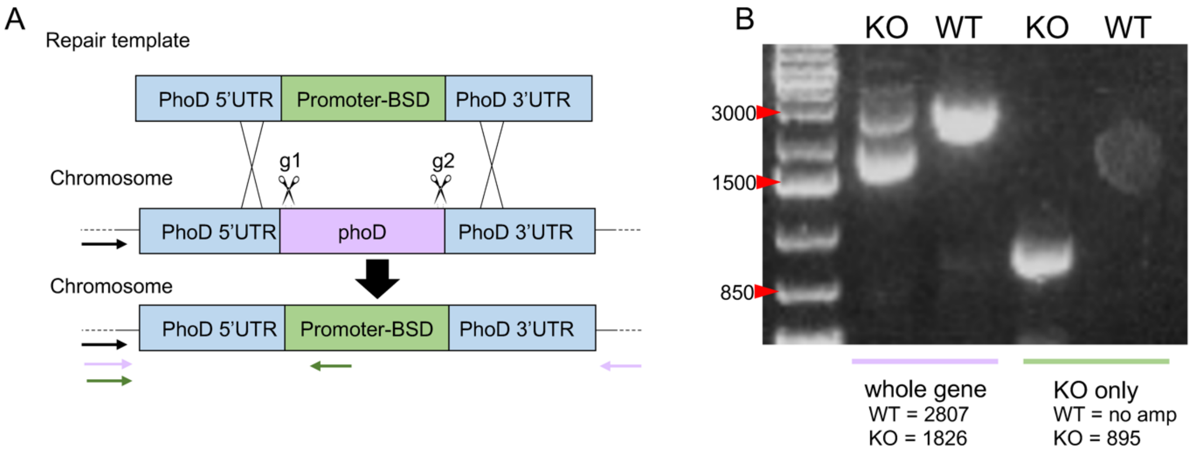


**SI Figure 3. Knock out of PhoD was unsuccessful.** (A) Schematic representation of the CRISPR/Cas9 based strategy to KO BdPhoD. (B) Gel image showing the diagnostic PCR to assess knock out. Lane 1 = Ladder (1 kb+), Lane 2 = Whole locus PCR for KO clones, Lane 3 = Whole locus PCR for WT clones, Lane 4 = KO specific PCR for KO clones, Lane 5 = KO specific PCR for WT. Primer locations for the whole locus PCR are shown on panel A as green arrows; primer locations for the KO specific PCR are shown on panel A in lavender arrows.

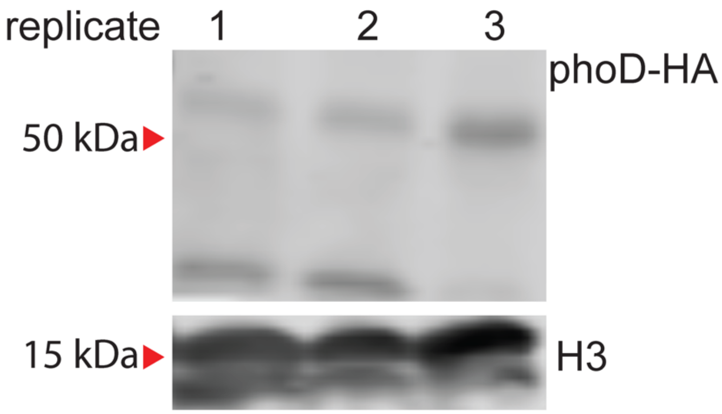


**SI Figure 4.** Confirmation of protein expression in overexpression parasites was done by immunoblotting for α-HA. 3 biological replicates are shown. α-H3 is used as the loading control.

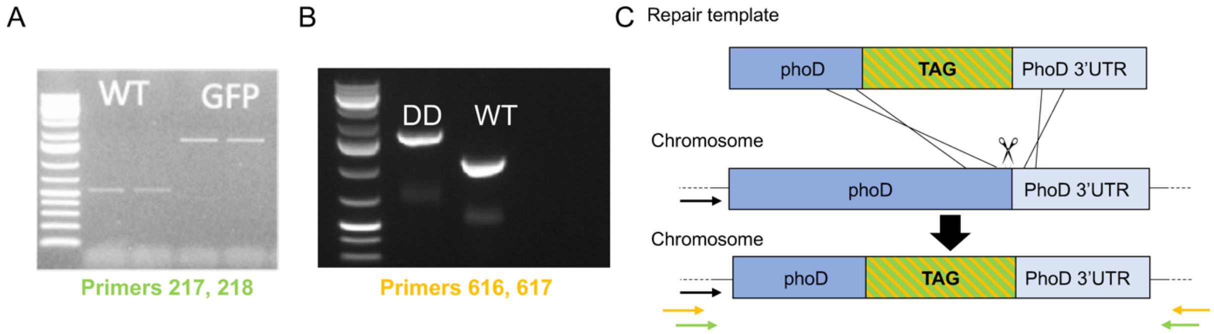


**SI Figure 5. Confirmation of endogenous tagging of BdPhoD (GFP, DD).** (A) Confirmation of integration of the 3' DD tag at the BdPhoD locus. Primers are designed to be outside of the homology repair template. Integration of the tag leads to an increase in band size. Lane 2, 3 = WT; Lane 4, 5 = PhoD-GFP. (B) Confirmation of integration of the 3' GFP tag at the BdPhoD locus. Primers are designed to be outside of the homology repair template. Integration of the tag leads to an increase in band size. Lane 2= PhoD-DD; Lane 3 = WT. (C) Schematic of the tagging strategy. The locations of the diagnostic primers are represented in green (GFP) and orange (DD). The primer sequences can be found in SI Table 1.

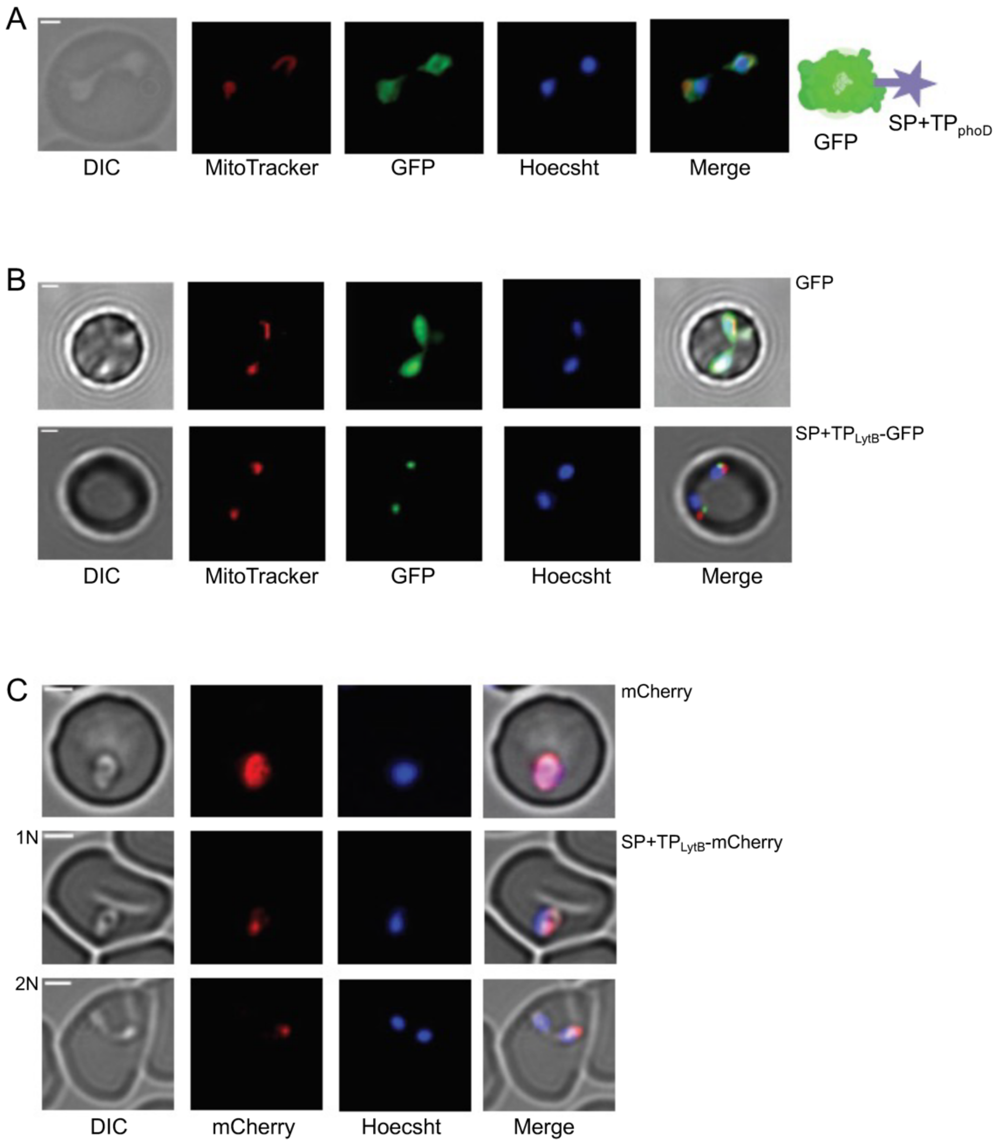


**SI Figure 6. Controls for IFA experiments.** In all images, Scale bar represents 2 μm. (A) Live cell images of *B. divergens* expressing ectopic GFP with the signal and transit peptides of phoD fused to the N-terminus, stained with MitoTracker$^{TM}$ Red. The localization is the same as the endogenous GFP tag on the C-terminus of PhoD. Schematic to the right represents the protein expressed from the construct. (B) Live cell images of *B. divergens* stained with MitoTracker$^{TM}$ Red. Top shows ectopic, cytoplasmic GFP expression. Addition of the signal peptide and leader peptide of LytB to GFP (pBd-SP+TP$_{lytB}$-GFP) results in apicoplast specific localization (bottom). (C) mCherry behaves the same as GFP in *B. divergens*. Top shows cytoplasmic mCherry. Middle- SP+TP$_{lytB}$-mCherry localizes to the apicoplast in both 1N (middle) and 2N (bottom) parasites.

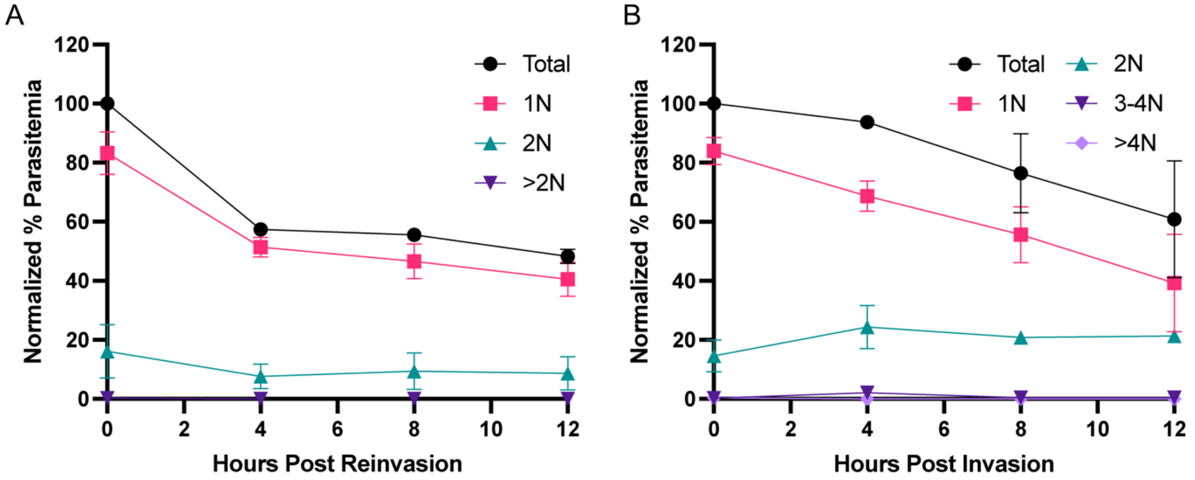


**SI Figure 7. MMV019266 inhibits division in *B. divergens* and *B. bovis*.** (A) Plot of flow cytometry data measuring nuclear content over time as a proxy for parasites/cell (hours post invasion) of synchronous *B. bovis.* Parasitemia is normalized to the starting parasitemia at time 0 h (n = 3). (B) Plot of flow cytometry data measuring nuclear content over time (hours post invasion) of synchronous *B. divergens*. Parasitemia is normalized to the starting parasitemia at time 0 h (n = 3).

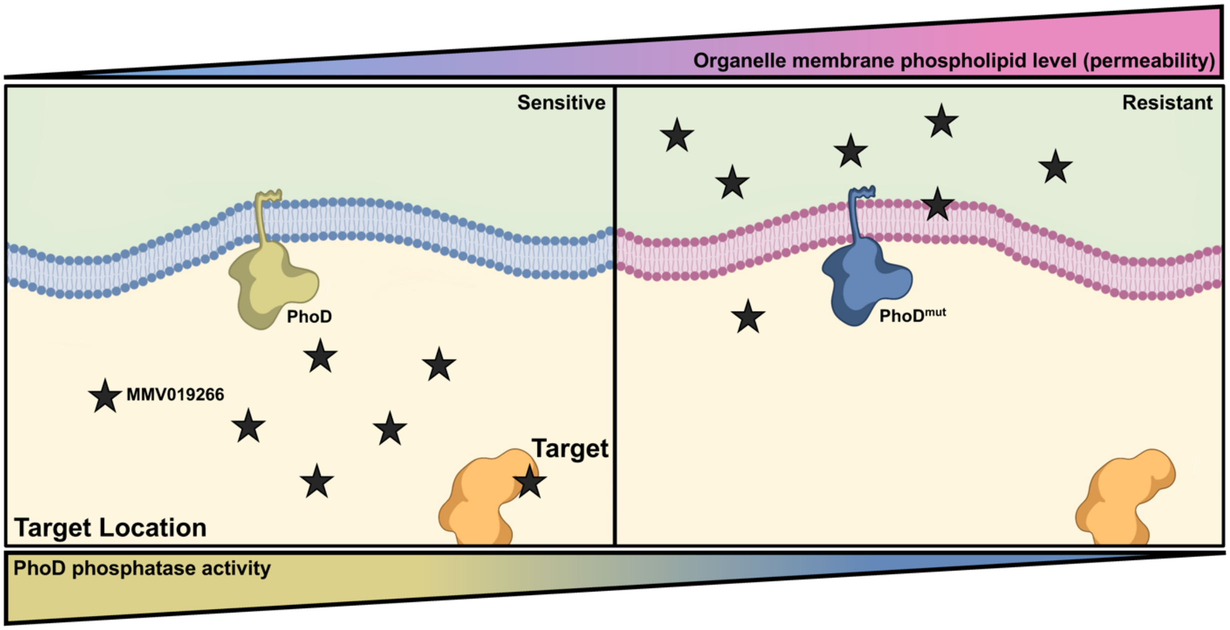


**SI Figure 8.** In this model, phoD is embedded in the membrane of the ER/apicoplast (endomembrane system). The catalytic domain points towards the target location (either the cytoplasm or the organelle lumen) and dephosphoylates the membrane leaflet surrounding it, altering the permeability of the membrane to MMV019266. The left represents the sensitive condition, where phoD activity is normal, allowing MMV019266 to access its target. The right panel demonstrates the resistant state, where the catalytic activity of phoD is decreased, altering membrane permeability, allowing for sequestration of MMV019266.
