## Supplemental Information for "Comparative chemical genomics in *Babesia* species identifies the alkaline phosphatase phoD as a novel determinant of resistance"

**SUPPLEMENTARY INFORMATION**

**Copy number variation analysis**

Read depth was assessed by plotting the reads aligned at each genome position from the BAM files. Depth was assessed looking at the ratio of depth between the parental strain and sample strain in 5 kb windows. Windows with very low coverage were excluded from the analysis.

*Babesia divergens:*

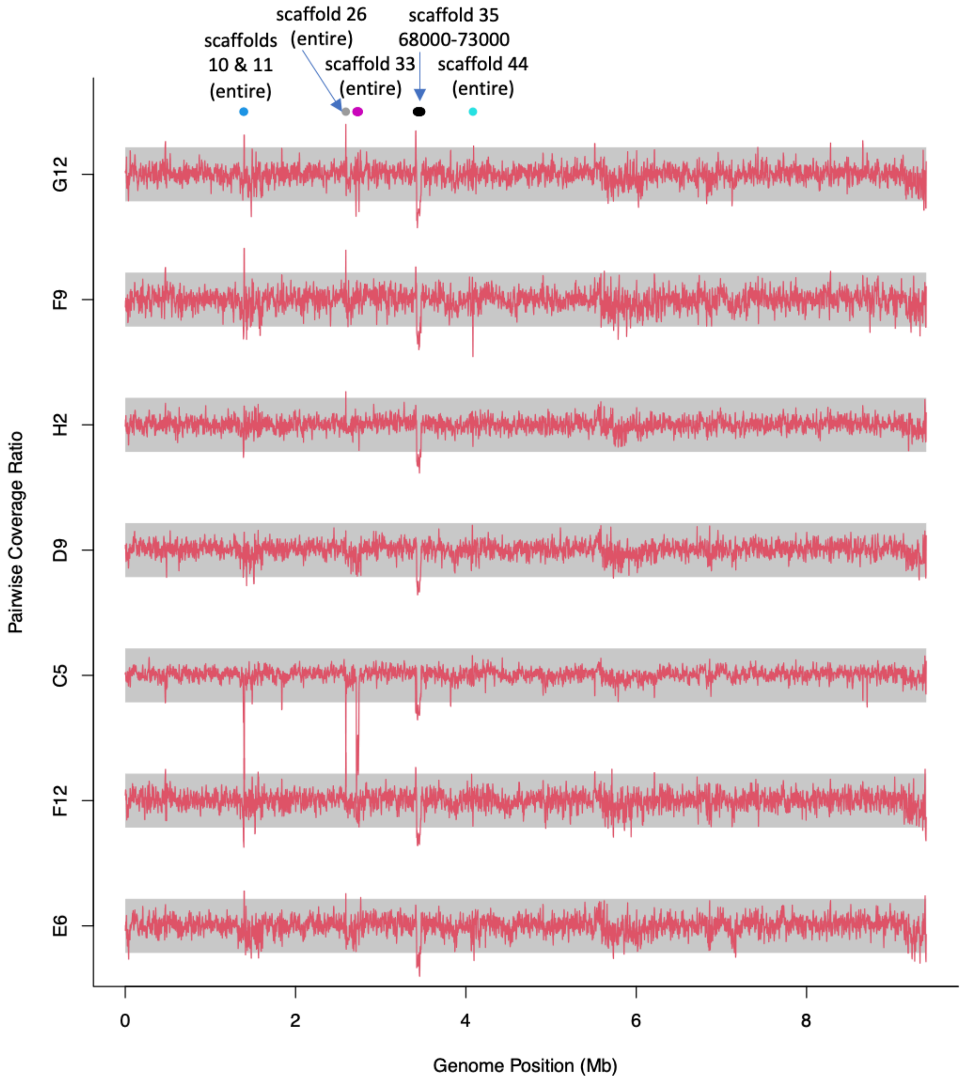


**Figure 1.** Log-scale coverage ratio between each sample and parental sample in 5-kb windows (windows with very low overall coverage excluded). Grey bars are the bounds of a 2-fold difference (lower edge = 0.5, upper edge = 2). Potential CNV sites are shows with colored dots and labeled for their genomic annotation.

In *Babesia divergens*, several potential regions of increased depth were identified (Fig 1). In each case, these regions either contain no gene, or a surface antigen VAR gene- neither of which are likely associated with resistance. The genome positions with increased depth in scaffold 10, 11, and 26, there are no genes in the enriched locations (Fig 2 A-C). For scaffold 44, all enriched regions correspond to VAR genes (Fig 2 D). These scaffolds are also very small contigs.

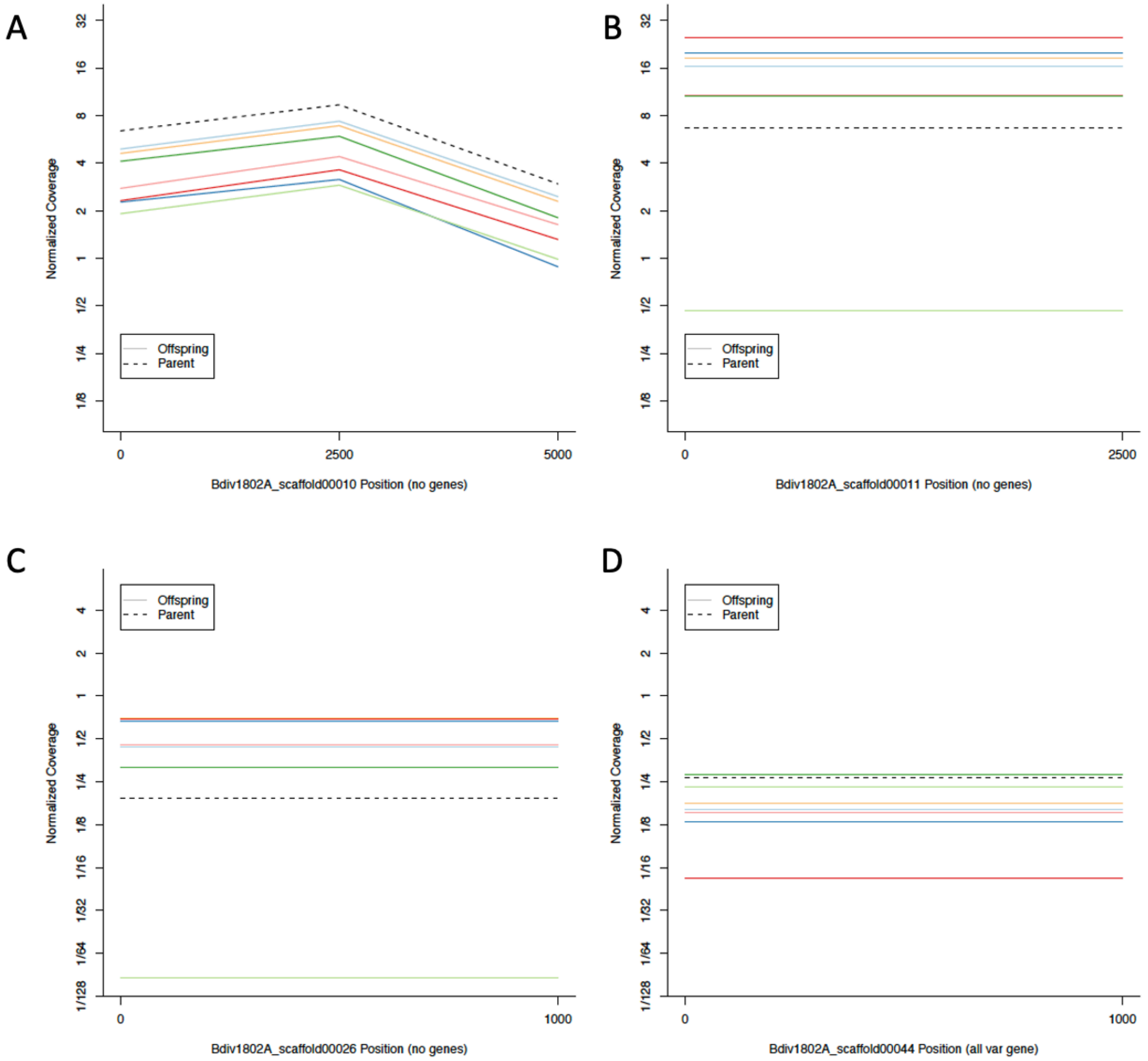


**Figure 2.** Coverage of specific genomic regions identified in the total analysis (5 kb windows). Each color represents a separate clone and the parental line is represented by a dashed line. Coverage per site is shown on the y-axis, with genome location on the x-axis. A) scaffold 10. B) scaffold 11. C) scaffold 26. D) scaffold 44.

Interestingly, in the parental strain it appears that a large segment of scaffold 35 is duplicated: however this may be due to poor genome annotation (Fig 3A). Further, there is very low coverage of scaffold 33 in clone C5 (Fig 3B).

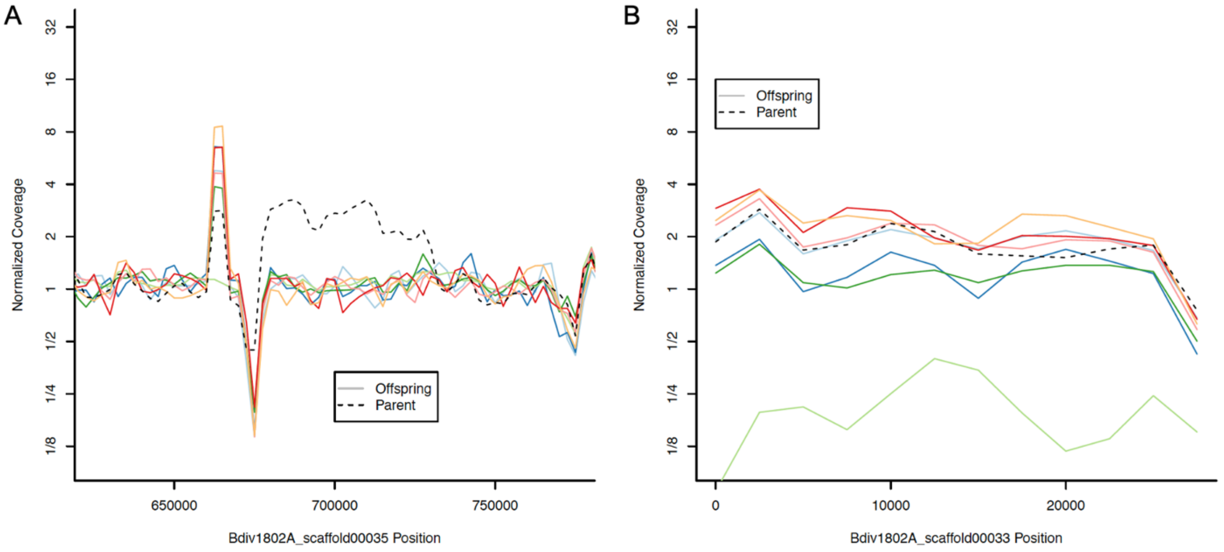


**Figure 3.** Coverage of specific genomic regions identified in the total analysis (5 kb windows). Each color represents a separate clone and the parental line is represented by a dashed line. Coverage per site is shown on the y-axis, with genome location on the x-axis. A) scaffold 35. B) scaffold 33.

Finally, we found no evidence of enrichment at the genomic region where BdPhoD is located (Fig 4).

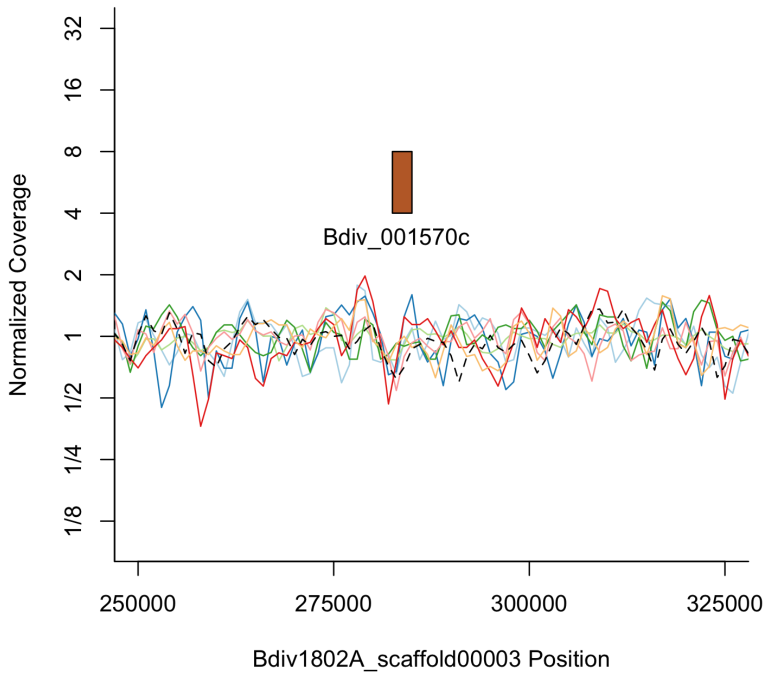


**Figure 4.** Coverage of specific genomic regions identified in the total analysis (2 kb windows). Each color represents a separate clone and the parental line is represented by a dashed line. Coverage per site is shown on the y-axis, with genome location on the x-axis. BdPhoD (Bdiv_001570c) is denoted by the brown box.

Taken together, these analyses reveal no compelling of CNVs contributing to resistance in *B. divergens*

B. bovis:

We plotted coverage per sample versus WT. Unexpectedly, there are many differences all samples share, which may indicate poor genome alignment, or incorrect parental strain (we identified this with several different sequencing preparations of the parental strain and observe the same pattern – data not shown). Regardless, we noted the large proliferation of mtDNA (far right) (Fig 5).

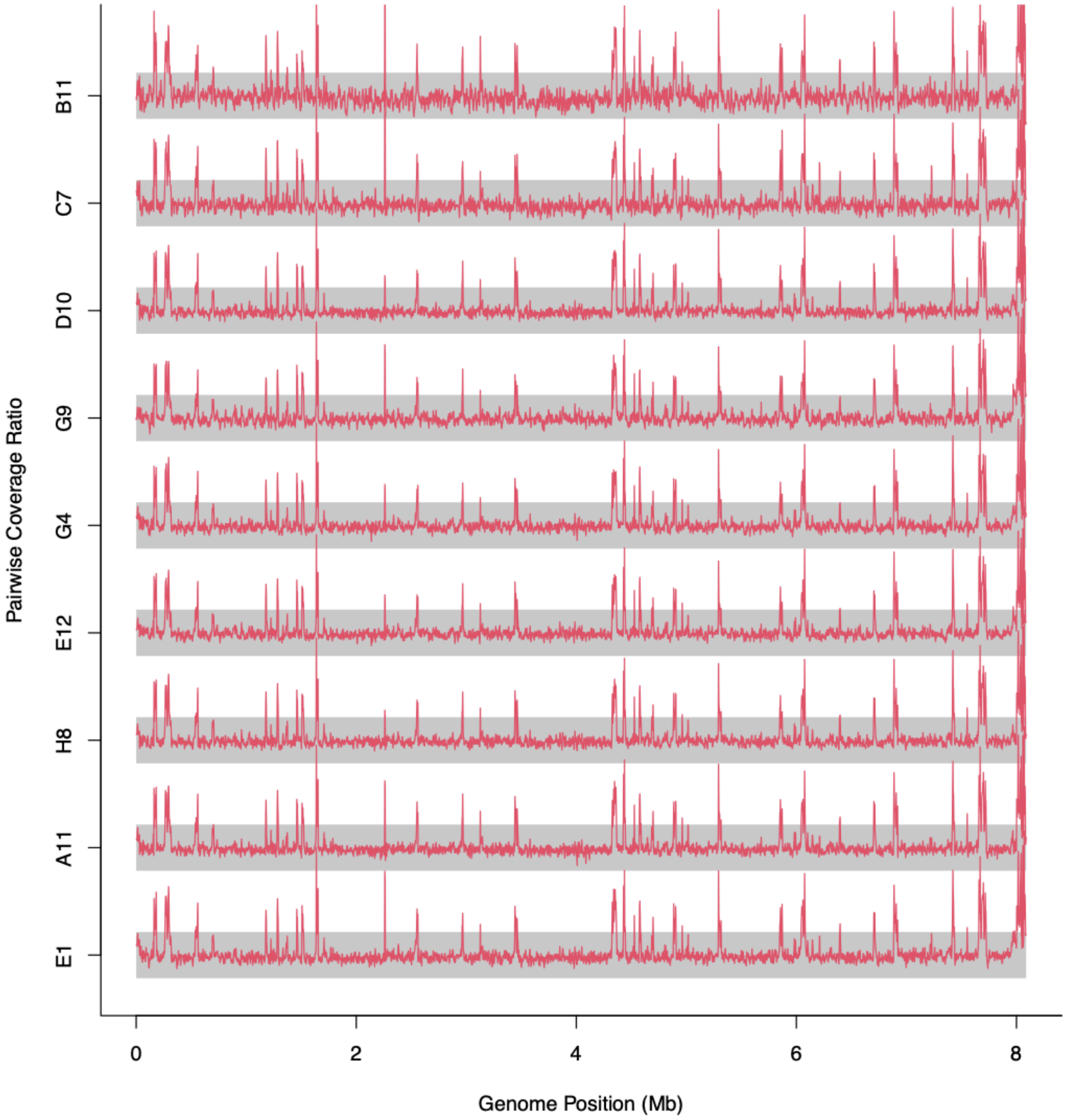


**Figure 5.** Log-scale coverage ratio between each sample and parental sample in 5-kb windows (windows with very low overall coverage excluded). Grey bars are the bounds of a 2-fold difference (lower edge = 0.5, upper edge = 2).

To overcome this noise, we instead performed CNV analysis comparing each clone to the average of all offspring (clone) samples). In this analysis we observed the same outlier as in Fig 5 (Fig 6).

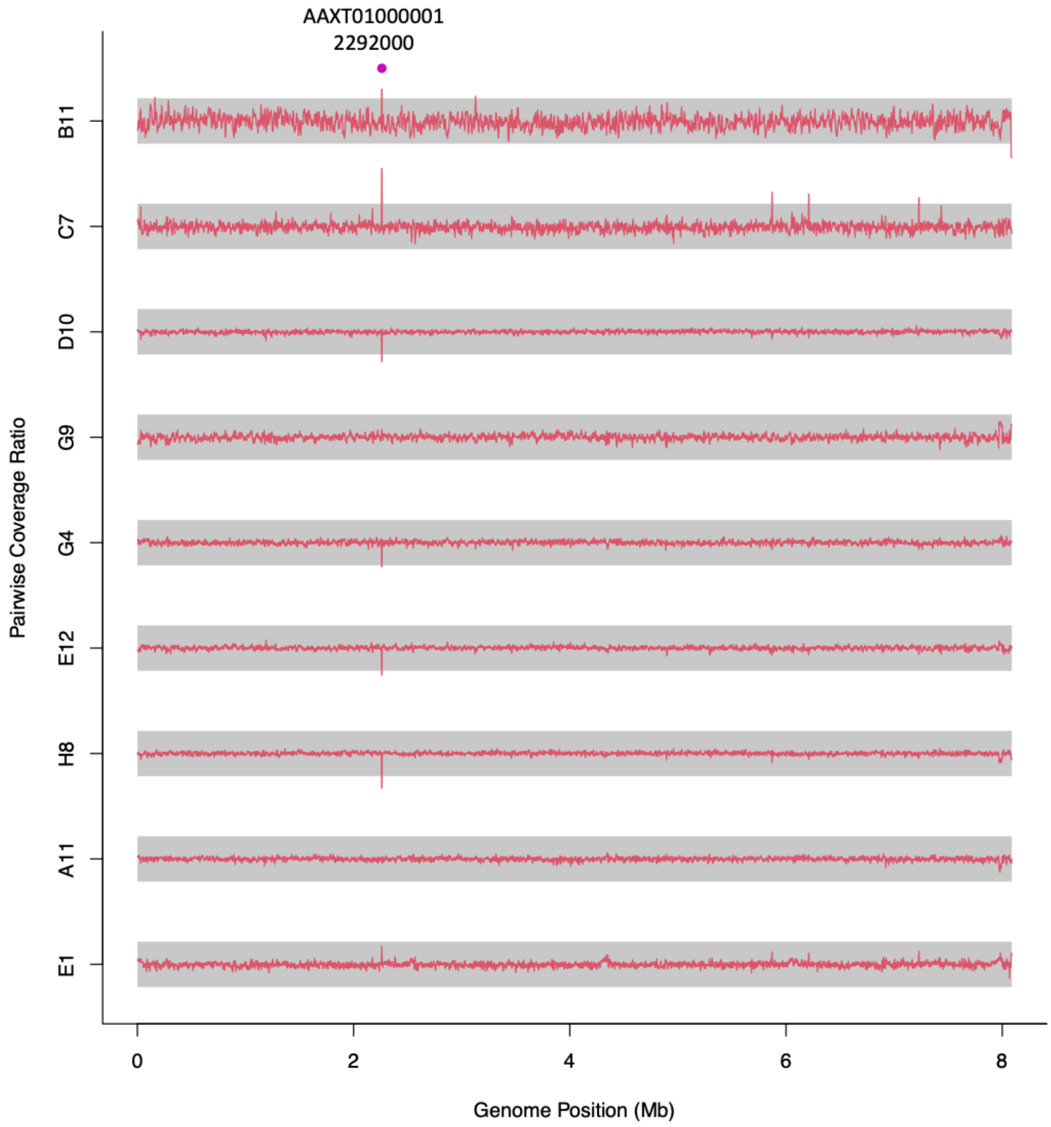


**Figure 6.** Log-scale coverage ratio between each sample and parental sample in 5-kb windows (windows with very low overall coverage excluded). Grey bars are the bounds of a 2-fold difference (lower edge = 0.5, upper edge = 2). Potential CNV sites are shows with colored dots and labeled for their genomic annotation.

The amplified region corresponds to the gene BBOV_III010730. This gene is a predicted Paf1 family protein. We observed amplification of this gene in selections with other, unrelated compounds (diminazene aceturate- data not shown), and conclude this is likely related to multidrug resistance, or a stress response for the parasite. Additionally, this gene was not identified as amplified in *B. divergens* and was observed in all clones of *B. bovis*- suggesting this may have been a hitch-hiking mutation unrelated to resistance.

Finally, we found no evidence of CNV at the genomic region where BbPhoD resides (Fig 7).

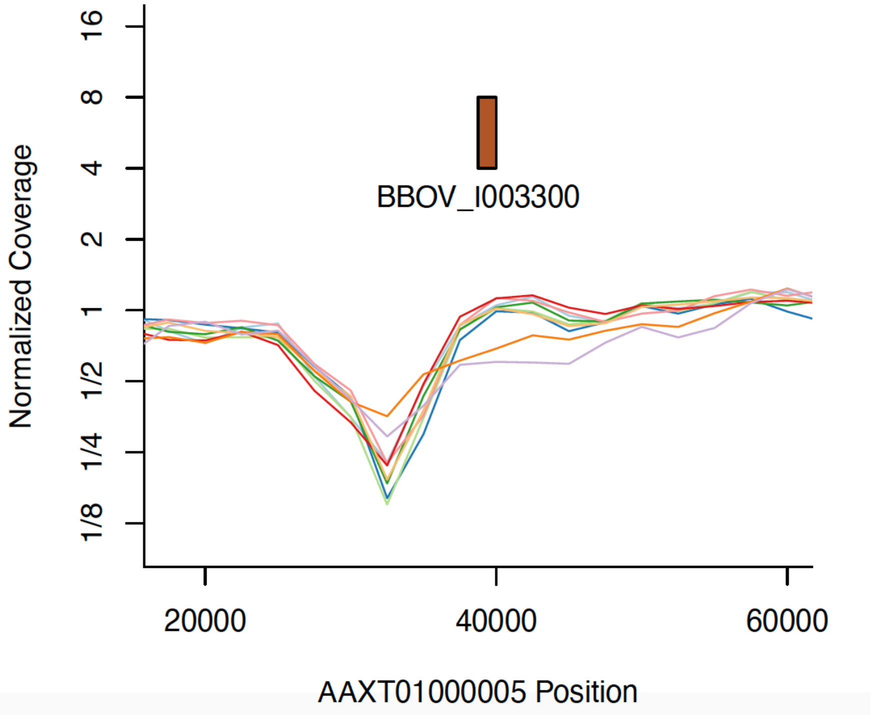


**Figure 7.** Coverage of specific genomic regions identified in the total analysis (2 kb windows). Each color represents a separate clone and the parental line is represented by a dashed line. Coverage per site is shown on the y-axis, with genome location on the x-axis. BbPhoD (Bdiv_I003300) is denoted by the brown box.

In conclusion, we found no compelling evidence of CNV contributing to resistance specific to MMV019266 in a species transcendent fashion.
