## Supplemental Table 1 for "Comparative chemical genomics in *Babesia* species identifies the alkaline phosphatase phoD as a novel determinant of resistance"

**SI Table 1. Sequences of primers used**

| **Primer name** | **Primer sequence (5'-3')** |
| --- | --- |
| PhoD_promoter_F | tgttgagcgtTATTTTGTTCATTCTGTAAGTATCC |
| PhoD_promoter_R | acggcaccatCATTCCGTTTGATTGGCAATC |
| PhoD_gDNA_F | aaacggaatgATGGTGCCGTATTGGTTATG |
| PhoD_gDNA_R | cgtaagggtaGTGCGTTTTCTCTTTGTG |
| 5' PhoD_GFPtag_F | tgggatatattagccgtatc*ctag*GTGTGCTGAAGTGGGAGGTAA |
| 5' PhoD_GFPtag_R | caccgccaccGTGCGTTTTCTCTTTGTGTTG |
| GFPtag_F | gaaaacgcacGGTGGCGGTGGCTCGATGGTGAGCAAGGGCGAGGA |
| GFPtag_R | ctgaaaaaacctacttgtacCTACTTGTACAGCTCGTCCATGCC |
| 3' PhoD_GFPtag_F | gtacaagtagGTTTTTTCAGTTGCTATATTTCTAC |
| 3' PhoD_GFPtag_R | ctcactatagaattcttaat*taa*GGAGTTTCCCACAGAACCTCCAGAC |
| KD_tag_F | ACCGGTTACCCTTACGATGTTCCTGACTATGC |
| KD_tag_R | gtcgacAGATCATGTGATTTCTCTTTGTTCAAGG |
| GFP_tag_check_F | TCTCGGGGGATGTTCATTACG |
| GFP_tag_check_R | GTTGACTTGGGGGCTACCAC |
| 5' PhoD_KD_HR1_F | tgggatatattagccgtatcctaggcgcagcagcagtcacaatactttacagG |
| 5' PhoD_KD_HR1_R | acatcgtaagggtaaccggtGTGCGTTTTCTCTTTGTGTTGACGTG |
| 3' PhoD_KD_HR1_F | gaaatcacatgatctgtcgacGTTTTTTCAGTTGCTATATTTCTACGTCGG |
| 3' PhoD_KD_HR1_R | ctcactatagaattcttaattaaGTAGAGGAATACCTTGTTCGCTTGG |
| KD_Tag check_F | TCTCGGGGGATGTTCATTACG |
| KD_Tag_check_R | GTTGACTTGGGGGCTACCAC |
| 5'_C196W_phoD_F | GGCCAATAGTAAAAACAGGATCTGTGG |
| 5'_C196W_phoD_R | tgggatatattagccgtatcctagGAAGCCATATACGATGACCATG |
| 3'_C196W_phoD_F | CCTGTTTTTACTATTGGCCTTGTC |
| 3'_C196W_phoD_R | ctcactatagaattcttaattaaGATATCAAATTGGCTATGCG |
| SNP_check_F | AGGCCTTTCTCTTGCAGCAT |
| SNP_check_R | TGTGTGCGTTATATCTGTCGCT |
| KD_Guide_Primer_F | CAAGTATCGATACAGGGAGGTTATCTGCTGAACAAGGTTACAgttttagagctaGAAA  tagcaagttaaaataagg |
| KD_Guide_Primer_R | CCTAGGATACGGCTAATATATCCCATC |
| GFP_guide_oligo_F | ggttATCTGCTGAACAAGGTTACA |
| GFP_guide_oligo_R | aaacTGTAACCTTGTTCAGCAGAT |
| SNP_guide_oligo_F | ggttTCCACAGGCATGTTTGACAA |
| SNP_guide_oligo_R | aaacTTGTCAAACATGCCTGTGGA |
| 5' PhoD_KO_F | tgggatatattagccgtatc*cta*GGATACCGACGAACAACG |
| 5' PhoD_KO_R | ctcactatagaattcttaat*taa*GTCATTTTGAAGATTTGTGTTGAC |
| KO_check_whole_F | GATGTATTCATAGATGCCGTCTAATGCG |
| KO_check_whole-R | CGTTTCGCACGCCATGCCATTCGCTCCTC |
| KO_specific_check_F | GTTAATGAGGATTCGAAATACGCTAC |
| KO_specific_check_R | CACGCTTAACGACAACGGGCACGAAG |
| KO_guide1_F | ggttGTGCCGTATTGGTTATGTGC |
| KO_guide1_R | aaacGCACATAACCAATACGGCAC |
| KO_guide2_F | ggttACACAAAGAGAAAACGCACT |
| KO_guide2_R | aaacAGTGCGTTTTCTCTTTGTGT |
